# Regulation of mRNA translation by a photoriboswitch

**DOI:** 10.1101/761775

**Authors:** Kelly A. Rotstan, Michael M. Abdelsayed, Luiz F. M. Passalacqua, Fabio Chizzolini, Kasireddy Sudarshan, A. Richard Chamberlin, Jiří Míšek, Andrej Lupták

**Affiliations:** Department of Pharmaceutical Sciences, University of California, Irvine, 92697, USA.; Department of Molecular Biology and Biochemistry, University of California, Irvine, 92697, USA.; Department of Chemistry, University of California, Irvine, 92697, USA.; Department of Organic Chemistry, Charles University, Prague, Czech Republic.

## Abstract

Optogenetic tools have revolutionized the study of receptor-mediated biological processes, but such tools are lacking for the study of RNA-controlled systems. To fill this gap, we used *in vitro* selection to isolate a novel RNA that selectively binds the *trans* isoform of a stiff-stilbene (amino-*t*SS), a rapidly and reversibly photoisomerizing small molecule. Structural probing revealed that the RNA binds amino-*t*SS about 100-times stronger than amino-*c*SS, giving the system robust selectivity for the *trans* isomer. *In vitro* and *in vivo* functional analysis showed that the riboswitch, termed Werewolf-1 (Were-1), inhibits translation of a downstream open reading frame when bound to amino-*t*SS and photoisomerization of the ligand with a sub-millisecond pulse of light induced the protein expression. Similarly, bacterial culture containing the *cis* isoform (amino-*c*SS) supported protein expression, which was inhibited upon photoisomerization to amino-*t*SS. Reversible regulation of gene expression using a genetically encoded light-responsive RNA will broaden the analysis of complex RNA processes in living cells.

---

Optogenetic techniques have transformed the biomedical sciences by controlling biological events with high spatiotemporal resolution through triggering signal transduction pathways *via* light-sensing proteins ^1–4;^ however, there are currently no photoactive molecules that can reversibly regulate cellular events at the RNA level. Photo-caged ligands have previously been used to regulate RNA, but their photo-uncaging procedures require relatively long UV light exposure and are irreversible, allowing only a single molecular event to be initiated ^5–10^. Furthermore, a light-responsive ribozyme has shown reversible activity, ^11^ and aptamers that bind photo-reversible ligands have been identified ^12–14^, but these RNAs have only been used *in vitro*. Here we used *in vitro* selection ^15–17^ to isolate a novel RNA that selectively binds only one photoisoform of a ligand, amino *trans* stiff-stilbene (amino-*t*SS) ^18–20^. Chemical probing identified amino-*t*SS–induced RNA structural changes in both the aptamer domain and a downstream expression platform derived from a bacterial riboswitch. *In vitro* and *in vivo* functional analysis showed that the riboswitch, termed Were-1, can induce or inhibit translation of a downstream open reading frame upon exposure to a sub-millisecond pulse of light, through reversible photoisomerization of the ligand. Our results demonstrate how a genetically encoded light-responsive RNA can reversibly regulate gene expression using light, providing a new optogenetic tool to broaden the analysis of complex RNA processes in living cells ^20–23^.

To isolate a new aptamer fused to a functional expression platform, we constructed an RNA pool derived from a bacterial SAM-I riboswitch ^24^ by replacing its ligand-binding domain with a 45-nucleotide random sequence, partially randomizing its anti-terminator and terminator hairpins, and retaining its translation initiation sequences (Supplementary Fig. 1). We synthesized a photoactive ligand – a *trans* stiff stilbene with an amino-terminated linker (amino-*t*SS) designed to maintain good cell permeability (Fig. 1a). The ligand was also designed to have a narrow window for photoregulation of both isomerizations in order to keep the rest of the visible spectrum available for the potential readouts of luminescent assays. Amino-*t*SS was characterized using UV-Vis and NMR spectroscopy to confirm photoisomerization to the *cis* conformation at 342 nm and back to the *trans* conformation at 372 nm, and to ensure that both isoforms are stable on timescales relevant to pulsed gene expression (Supplementary Fig. 2). The RNA pool was selected *in vitro* to bind amino-*t*SS coupled to carboxylate agarose beads and eluted under denaturing conditions ^16^. We hypothesized that a pool of amino-*t*SS–binding aptamers would include motifs that do not bind the *cis* photoisoform of the ligand and that the *t*SS–binding conformation stabilizes the expression platform in a single state that affects either transcription or translation of a downstream open reading frame (ORF). After six rounds of *in vitro* selection, we cloned the pool into bacterial plasmids and tested individual sequences for amino-*t*SS binding by monitoring RNA–dependent changes in the fluorescence of the amino-*t*SS. One sequence showed markedly increased fluorescence of amino-*t*SS (Supplementary Fig. 3, Fig. 1c). This sequence, termed Werewolf-1 (Were-1) for its potential light-dependent conformational changes, was chosen for further analysis.

**Figure 1.**
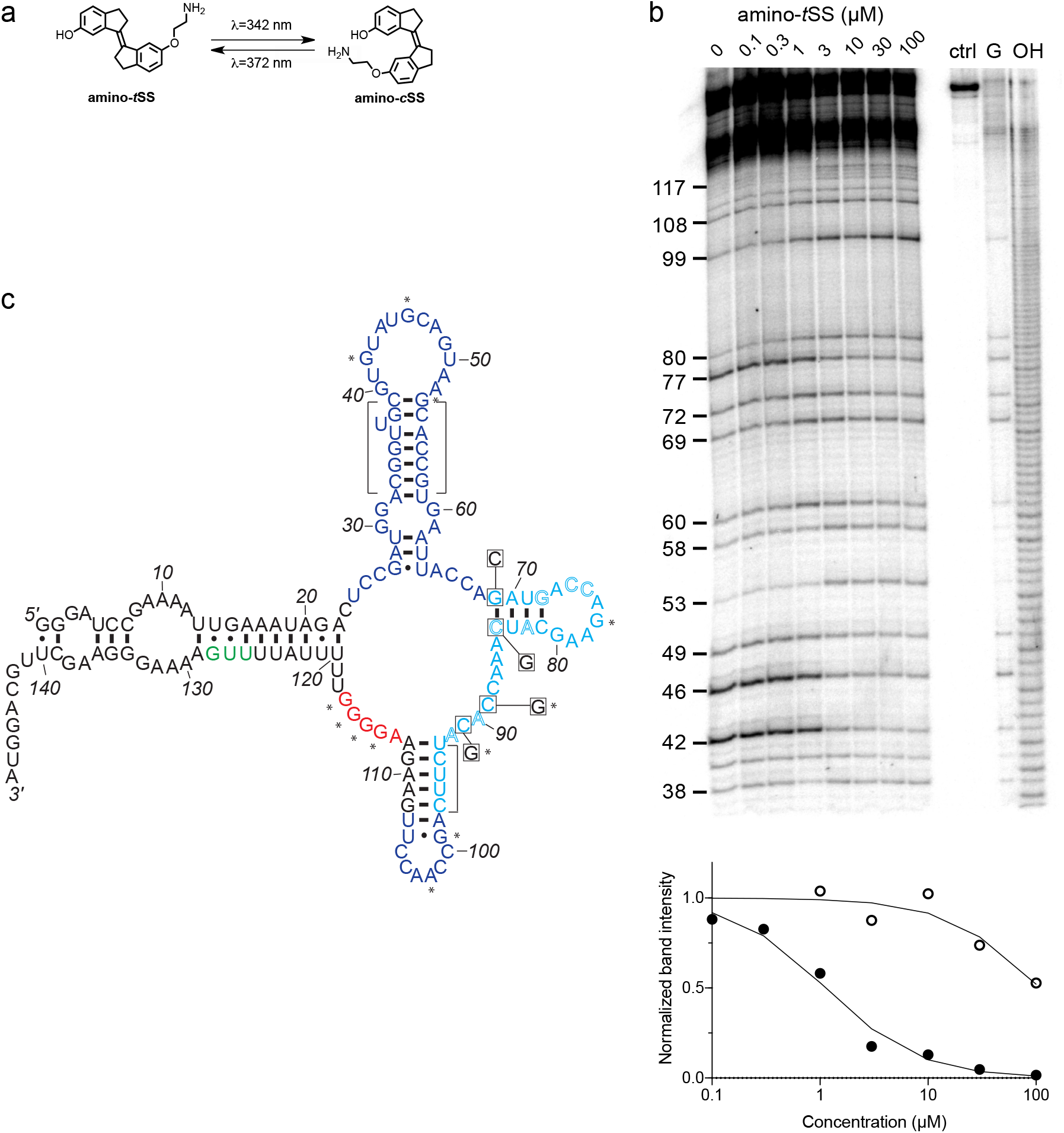
An amino-*t*SS-responsive aptamer. **a**, Amino-*t*SS isomerizes from *trans* to *cis* conformation when exposed to 342 nm light, and back to the *trans* isoform at 372 nm. **b**, RNase T1 probing of Were-1 structure. Right lanes contain a control with undigested RNA (ctrl), a T1-digested sequencing control (G), and a hydroxide-mediated partial digestion ladder (OH) of the RNA. The left lanes show partial T1 digestion in the presence of increasing amino-*t*SS at concentrations indicated above the gel image. The probing shows clear ligand-dependent changes—both increases (e.g. G53, G99, G114-117) and decreases (e.g. G 42, G46, G77)— interspersed throughout the sequence. Below, an apparent *K*_D_ of 1.1 *µ*M was calculated based on the change in band intensity with increasing amino-*t*SS (dark, filled circles) for nucleotide G46, normalized to a control band (G72). Additionally, a *K*_D_ of 108 *µ*M was calculated based on the change in band intensity with increasing amino-*c*SS (open circles) for the same nucleotide and control (Supplementary Fig. 4g), suggesting high specificity for amino-*t*SS. An average *K*_D_ value of 1.5 µM amino-*t*SS was calculated for changes in nucleotides G42, G46, G77, and G80. **c**, Secondary structure prediction of Were-1 derived from all structural probing data in absence of the ligand (see also Supplementary Fig. 4). Partially randomized regions (light blue), the Shine-Dalgarno sequence (red), the start codon (green), and the 3’ terminus sequence are derived from the *B. subtilis mswA* SAM-I riboswitch. The 5’ part of the aptamer (dark blue) was selected from the random region of the starting pool (Fig. S1). Outlined letters are positions where the selected sequence differs from the *B. subtilis* riboswitch expression platform. Boxed positions were mutated to the indicated nucleotides to identify regions of structural and functional importance. Bracketed regions indicate areas that do not change in the presence of amino-*t*SS, and asterisks (*) indicate nucleotide positions that do change in the presence of amino-*t*SS.

To assess the ligand-dependent structural modulation of Were-1, we performed multiple RNA structure-probing experiments, including digestions with T1 and S1 nucleases ^25, 26^, terbium (III) footprinting ^27^, in-line probing ^28^, and selective 2’ hydroxyl acylation by primer extension (SHAPE) ^29^ using a range of amino-*t*SS concentrations (Fig. 1b, Supplementary Fig. 4). The changes in the pattern of RNA probing suggested that Were-1 undergoes conformational modulation upon introduction of amino-*t*SS in both the sequence derived from the randomized region and the expression platform (Fig. 1b, Supplementary Figs. 4a–4c, and 4g). Control experiments with amino-*t*SS analogs, such as *trans*-stilbene (*t*S), 4,4-*trans*-dihydroxystilbene (*t*DHS), and S-adenosyl methionine (SAM), showed no change in the probing patterns (Supplementary Figs. 4c–4f), suggesting that Were-1 is specific for amino-*t*SS. Analysis of the T1 probing experiments revealed a *K*_D_ of 1.1 *µ*M, based on the change in band intensity with increasing amino-*t*SS for nucleotide G42, normalized to a control band G72 (Figure 1b). Additionally, a 108-µM *K*_D_ was calculated based on the same band intensity change with increasing amino-*c*SS (Supplementary Fig. 4g), revealing a 100-fold specificity for the target ligand, amino-*t*SS. The average amino-*t*SS *K*_D_ derived form the T1 nuclease probing (at positions 42, 46, 77, and 80; normalized to the band intensity at position 72) was of 1.5 *µ*M, whereas the nuclease S1 probing revealed a *K*_D_ of 0.4 *µ*M (based on band intensity change for positions A44-G46), and the terbium (III) footprinting yielded a somewhat higher apparent *K*_D_ of 4.8 *µ*M (calculated based on the change in intensity at positions A113 and U107, normalized to G134 control band; Supplementary Fig. 4c). In order to establish the location of the amino-*t*SS aptamer domain, we modeled the secondary structure of Were-1 based on the probing data and created mutants hypothesized to affect ligand binding affinity or RNA structural stability (Fig. 1c) and tested them *in vitro* and *in vivo*. The secondary structure modeling did not support a conformation containing a Rho-independent transcriptional terminator, in part because the selected sequence contained two mutations (C90A and U92A), which are predicted to disrupt the stability of a full-length transcription-terminating helix (Fig. 1c).

To test whether Were-1 can directly couple light-induced states of the ligand to the activity of the expression platform *in vitro*, the aptamer was tested for amino-*t*SS–dependent conformational changes using a strand-displacement assay that mimics mRNA binding by the bacterial ribosome 17,30,31. We designed a DNA duplex in which the longer strand has a toehold sequence corresponding to the reverse complement of the Shine-Dalgarno sequence of Were-1 (Supplementary Table 1) and a fluorophore to assess whether the shorter DNA strand, containing a quencher chromophore, is displaced through RNA:DNA hybridization with Were-1 (Supplementary Fig. 5a). Polyacrylamide gel electrophoresis (PAGE)-purified Were-1 bound the toehold readily, but the strand displacement was diminished in the presence of amino-*t*SS (Supplementary Fig. 5b). Testing the toehold binding at various concentrations of amino-*t*SS revealed a dose dependence with a half–maximal inhibition of toehold binding at ∼6 *µ*M (Supplementary Fig. 5c).

We next asked whether the ribosome mimic binds this RNA during *in vitro* transcription (Fig. 2a, Supplementary Fig. 6). In the absence of the ligand, the RNA bound the toehold efficiently, showing a robust increase of fluorescence immediately after transcription initiation. In contrast, the addition of high concentration (14.8 *µ*M) of amino-*t*SS strongly abrogated new binding of the toehold, as revealed by almost a full reduction in the slope of the fluorescence expansion curve (Fig. 2b). Intermediate concentrations of amino-*t*SS were tested to assess RNA binding and specificity, yielding a ligand-dependent response with a half-maximum of ∼4 *µ*M amino-*t*SS (Fig. 2c). Probing-derived secondary structure modeling of Were-1 suggested substantial ligand-dependent conformational changes in multiple parts of the sequence, except for one predicted hairpin. To confirm the presence of this structural element, we created a variant containing a single mutation (G69C), that was predicted to disrupt this helix, and a presumed compensatory mutant (G69C/C84G) (Fig. 1c). The G69C variant showed diminished response to amino-*t*SS, whereas the G69C/C84G double-mutant exhibited partially restored activity, suggesting that these two positions are indeed part of a helix. Other variants, C89G and C91G, designed based on parts of the sequence that showed amino-*t*SS–dependent changes in the structure-probing experiments, both showed decreased sensitivity to the ligand, suggesting that they are essential for ligand binding. Furthermore, when testing toehold binding using the purified *cis* isoform of the stiff stilbene (amino-*c*SS), as well as other stilbenes, such as *t*S, *t*DHS, and *trans* stiff stilbene (*t*SS; Were-1 ligand lacking the aminolated linker), no significant changes in fluorescence were observed (Fig. 2c, Supplementary Fig. 7). These results demonstrate that amino-*t*SS stabilizes the RNA in an “OFF” (ribosome-inaccessible) conformation in a dose-dependent manner and with high ligand specificity.

**Figure 2.**
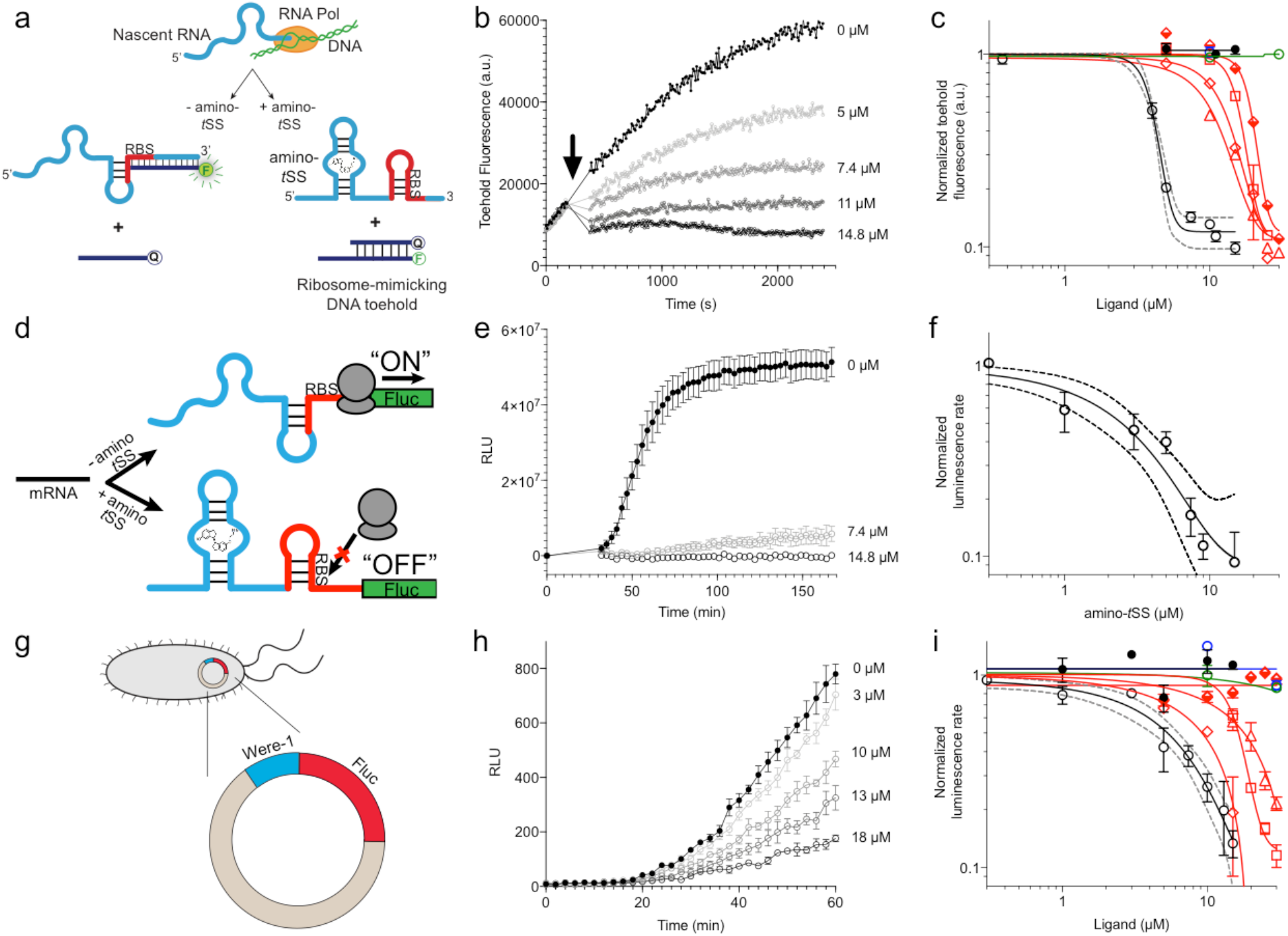
Translation regulation by the Were-1 riboswitch. **a**, Schematic of co-transcriptional binding of Were-1 RNA to amino-*t*SS in the presence of a toehold-reporter complex. In absence of amino-*t*SS, the transcribed RNA exposes the ribosomal binding site (RBS), enabling binding of the complementary region of the toehold reporter, displacing the quencher strand, and producing a fluorescence signal. In presence of amino-*t*SS, the RNA binds the ligand, sequestering the RBS and preventing displacement of the quencher strand. **b**, Co-transcriptional response of Were-1 to different concentrations of amino-*t*SS using the toehold reporter. Initial transcriptions of Were-1 without ligand show identical increase in toehold fluorescence for all samples. When amino-*t*SS is added (arrow), a dose-dependent decrease in fluorescence is observed. **c**, Response (± SEM; n=81) of Were-1 (black, open circles), and its variants (red) C89G (triangles), C91G (squares), G69C (half-shaded diamonds), and G69C/C84G (open diamonds), in the presence of amino-*t*SS shows a shift in dose-dependence for single mutations, particularly G69C, and partial recovery of activity for the G69C/C84G double mutant. Were-1 shows no response in the presence of amino-*c*SS (black circles), *trans*-stilbene (green, open circles) and *trans*-4,4-dihydroxystilbene (blue, open circles). **d**, Schematic of amino-*t*SS-dependent inhibition of protein expression *in vitro* using a Were-1-firefly luciferase (Were-1-Fluc) construct. In absence of the ligand, the RBS is exposed and luciferase is translated, whereas in presence of amino-*t*SS, the RBS is sequestered, abrogating Fluc expression. **e**, *In vitro* translation of the Were-1-Fluc construct. Robust luminescence is observed when no ligand is present, but the signal is significantly lower in presence of amino-*t*SS. **f**, Response (± SEM; n = 58) of the Were-1–regulated protein expression to amino-*t*SS. **g**, Schematic of the Were-1-Fluc construct incorporated into a bacterial plasmid. **h**, Were-1– controlled Fluc gene expression in *E. coli*. Bioluminescence is observed in absence of amino-*t*SS, and progressively diminished with increasing amino-*t*SS. **i**, Expression of Were-1-Fluc (± SEM; n=257) *in vivo* (black, open circles), and its variants (red) C89G (triangles), C91G (squares), and G69C/C84G (open diamonds), in the presence of amino-*t*SS, show a dose-dependent response. Were-1 mutant G69C (half-shaded diamond) and Were-1 in the presence of amino-*c*SS (black circles), *trans*-stilbene (green, open circles) and *trans*-4,4-dihydroxystilbene (blue, open circles) showed no change in bioluminescence, whereas the G69C/C84G double mutant shows restoration of activity similar to wild-type levels. Note, dose-response graphs (**c**, **f**, **i**) are on a log-log scale. The apparent amino-*t*SS IC_50_s are 3.9 ± 0.2, 2.5 ± 1.0, and 5.3 ± 1.1 µM for the toehold (**c**), *in vitro* translation (**f**), and *in vivo* expression (**i**), respectively. Dashed lines correspond to the 95% confidence interval of the binding model.

In order to test whether the Were-1 RNA interaction is selective for amino-*t*SS, and potentially acts as an amino-*t*SS riboswitch, we created a construct consisting of the putative riboswitch, including its minor start codon from *Bacillus subtilis* that was present in the starting pool, followed by a firefly luciferase (Fluc) ORF lacking its endogenous start codon. Based on the toehold assays, we hypothesized that in absence of amino-*t*SS, Were-1 would be transcribed in a conformation promoting the translation of the luciferase enzyme, whereas the presence of amino-*t*SS would stabilize a conformation preventing efficient translation initiation, downregulating the luciferase expression (Fig. 2d). Using a purified bacterial *in vitro* transcription/translation system, luciferase production was measured in the presence and absence of amino-*t*SS. In absence of the ligand, the construct exhibited robust luciferase production, demonstrating that the unbound aptamer promotes protein production from a downstream ORF (Fig. 2e). In contrast, when amino-*t*SS was added, protein production decreased in a dose-dependent manner (Fig. 2f). To confirm that the ligand itself did not impact the *in vitro* translation system, luminescence was tested with a control plasmid lacking the riboswitch, and no amino-*t*SS sensitivity was observed (Supplementary Fig. 8).

To further test the riboswitch, we incorporated the construct into a bacterial plasmid and induced its expression in *E. coli* cells (Fig. 2g). Bioluminescence, due to luciferase expression and activity, was robust in the absence of ligand, and when the cells were incubated in the presence of amino-*t*SS, bioluminescence was again diminished in a dose-dependent manner (Fig. 2h, i). To confirm the specificity of Were-1 for amino-*t*SS, we incubated cells in the presence of amino-*c*SS, *t*S and *t*DHS, and observed no significant change in bioluminescence (Fig. 2i). To determine the effect of an alternate start codon on Fluc expression, we changed the UUG start codon in the Were-1-Fluc plasmid to AUG. Bioluminescence was higher in the AUG samples compared to the wild-type UUG construct and showed a similar amino-*t*SS–dependent response (Supplementary Fig. 9). Testing the above-mentioned mutants confirmed the G69/C84 interaction, and the importance of the C89 and C91 positions for ligand binding. Taken together, our results demonstrate that Were-1 controls amino-*t*SS–dependent protein expression *in vitro* and *in vivo,* acting as a translational riboswitch.

We next asked whether Were-1 could regulate gene expression in a light-dependent manner, acting as a photoriboswitch. For this to occur, the *trans* isoform of the stiff stilbene must photoisomerize to its *cis* state, preventing binding of the Were-1 aptamer domain, and promoting expression of a downstream ORF.

Riboswitches are sensitive to co-transcriptional events because they are capable of adopting different RNA folds as they are transcribed by RNA polymerase ^32^. To first monitor changes in RNA folding over time, we used the toehold-fluorophore system to determine whether the DNA duplex was in a bound state (no strand displacement) versus an unbound state (strand displacement releasing the quencher DNA, yielding fluorescence) during transcription. Our results show that when the amino-*t*SS-bound Were-1 structure was irradiated at 342 nm of light, the toehold fluorescence increased, suggesting that the RNA increased the binding to the ribosome mimic present on the DNA toehold. Furthermore, when exciting amino-*c*SS at 372 nm of light to switch the ligand to its *trans* isoform, we observed a decrease in toehold fluorescence growth, indicating that the photo-generated amino-*t*SS was able to re-bind the Were-1 RNA. The irradiation was repeated until all toehold was bound, with each switch showing consistent results (Supplementary Fig. 10). This experiment shows reversible, wavelength-dependent binding of the ribosome mimic, emulating light-dependent protein expression from a downstream open reading frame.

To study the system further, we used the Were-1-luciferase construct and the purified bacterial *in vitro* transcription/translation system to test whether luciferase expression could be regulated by our putative photoriboswitch. We found that when the reaction was irradiated at 342 nm of light, luminescence increased, suggesting that Were-1’s conformation changed to expose the RBS, enabling luciferase expression. When the *cis* stilbene was photoisomerized to the *trans* state (amino-*t*SS) at 372 nm of light, luciferase protein production slightly decreased (Supplementary Fig. 11). These data are consistent with an *in vitro* activity of a photoriboswitch.

Finally, we used *E. coli* containing the Were-1-Fluc construct to determine whether gene expression can be regulated with a pulse of light. As shown above, bioluminescence was greatly diminished in the presence of amino-*t*SS (Fig. 2i). Upon exposing the bacteria to 342 nm light, we saw a robust increase in bioluminescence (Fig. 3a, b and Supplementary Fig. 12). This result strongly suggests that upon photoisomerization of amino-*t*SS to the *cis* isoform with a pulse of light, Were-1 was able to change conformation and expose its RBS to allow luciferase production. To confirm these results, we performed control experiments, in which cells were covered during the excitation to distinguish between regular *E. coli* growth behavior and the increase in bioluminescence from the riboswitch (Fig. 3a, b). Additionally, in another experiment, we excited cells with a different wavelength of light (500 nm) that should not impact the isomerization of the ligand (Supplementary Fig. 12). Both controls showed lower bioluminescence compared to the 342 nm excited samples. To further analyze the system, we tested the temporal dependence of Were-1–regulated Fluc production by exposing an amino-*t*SS–containing bacterial culture to 342 nm light for various lengths of time. Relative to controls that were unexposed, the highest luciferase expression was seen at an exposure time of 500 *µ*s (Fig. 3b). Furthermore, when testing the same system using amino-*c*SS, bioluminescence decreased in a dose-dependent manner with increasing amino-*c*SS concentration after exposure to 390 nm light (Fig. 3c). This result strongly suggests that by isomerizing amino-*c*SS to its *trans* isoform with a pulse of light, Were-1 was able to sequester its RBS to inhibit luciferase production. Testing the temporal response of Were-1–regulated Fluc expression in the presence of amino-*c*SS revealed that Were-1 regulates expression optimally at short exposures, showing the highest inhibition after a millisecond of light exposure (Fig. 3d). Based on isomerization data (Supplementary Fig. 2b), this effect is likely due to the ligand reaching a semi-photostationary state after longer light exposure. No difference in cell density was observed among the experiments, implying that neither the ligand, nor the light pulses affect the bacterial growth, and suggesting negligible photo-damage to the cells.

**Figure 3.**
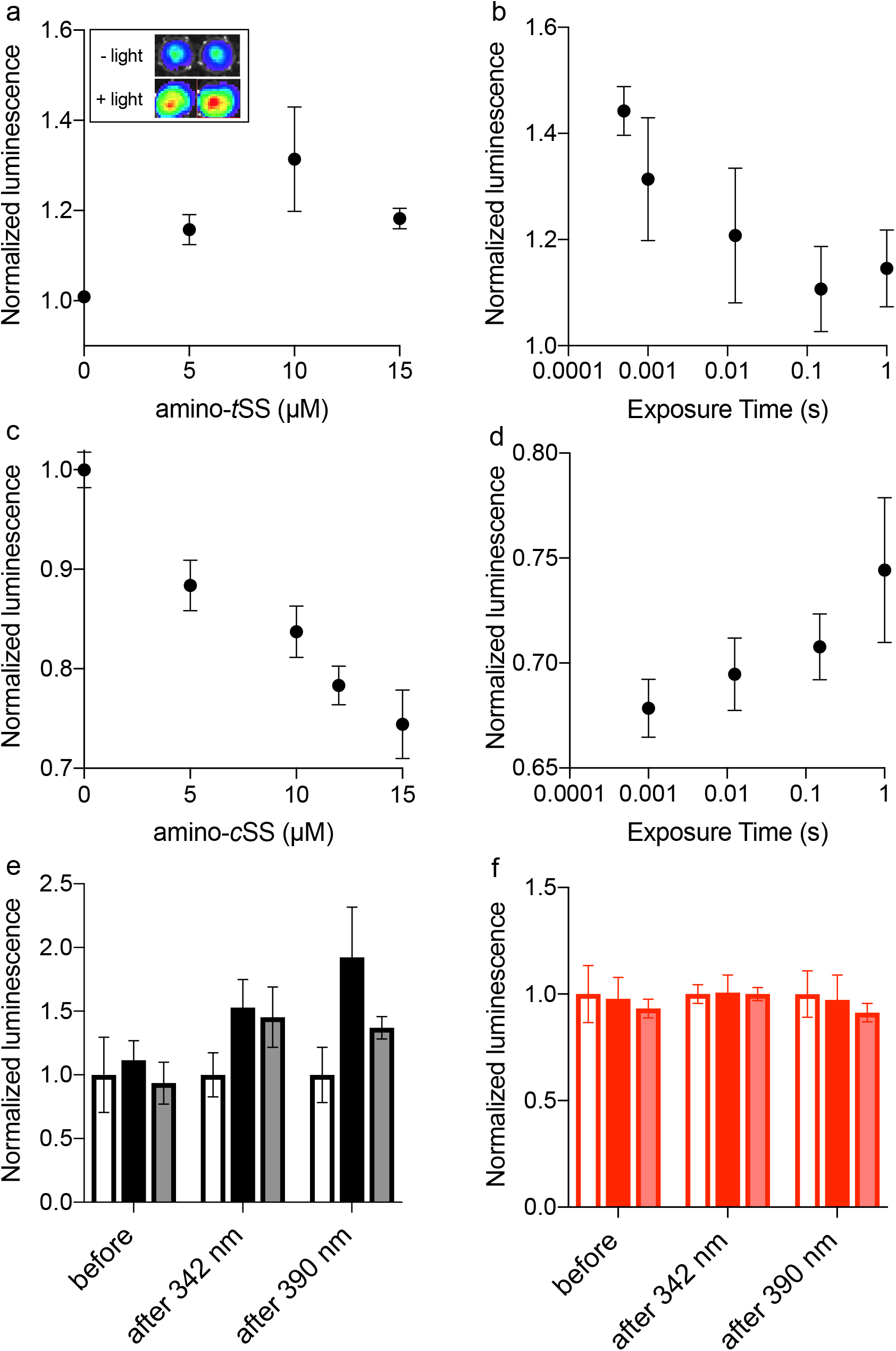
Regulation of luciferase expression by the Were-1 photoriboswitch *in vivo*. **a**, Normalized amino-*t*SS–dependent bioluminescence (± SEM) of the Were-1-Fluc construct after 1 ms exposure of 342 ± 5 nm light (Φ_q_ = 1.4*10^-2^ W/cm^2^). The largest change in expression was observed in the presence of 10 µM amino-*t*SS. Inset shows the light–dependent bioluminescence of the bacterial cultures at 10 µM ligand. **b**, Were-1 regulation of luciferase expression (± SEM) *in vivo* at various exposure times in presence of 10 µM amino-*t*SS. **c**, Normalized bioluminescence of the Were-1-Fluc *E. coli* incubated with amino-*c*SS after 1 s exposure of 390 ± 9 nm light (Φ_q_ = 5.5*10^-2^ W/cm^2^) showing progressively higher protein expression inhibition at higher amino-*c*SS concentrations, presumably due to higher concentration of amino-*t*SS after photoisomerization. **d**, Change of Fluc expression after photoisomerization of 15 µM amino-*c*SS at 390 ± 9 nm light for various exposure times, showing largest photoswitching at 1 ms exposure. **e,** Regulation of luciferase expression (± SEM) by the Were-1-Fluc construct before exposure, forty-five minutes after a 1-ms pulse of 342 ± 5 nm light (black and gray), which resulted in increased bioluminescence compared to control (clear), and forty-five minutes after 0.5 ms of 390 ± 9 nm exposure that resulted in decreased bioluminescence (gray) compared to samples that were only exposed to 342 nm (black). **f,** Luciferase expression (± SEM) by the Were-1-Fluc G69C mutant using the same conditions as above (e), showing no significant change in expression after exposure to 342 nm (red, light red) or 342 nm and 390 nm (light red) compared to unexposed controls (clear).

We next asked whether Were-1 could reversibly regulate gene expression *in vivo*, providing the first optogenetic tool to reversibly regulate cellular events at the RNA level. Using the same *E. coli* construct, we measured bioluminescence over two hours in samples that were covered during excitations, and therefore unexposed to light, samples that were exposed to a millisecond of 342 nm light, and samples that were exposed to 342 nm and then later excited at 390 nm for a sub-millisecond. Initial values prior to excitation showed no significant difference in bioluminescence twenty-five minutes post induction (Fig. 3e). Samples that were then exposed to 342-nm light showed a significant increase in luciferase expression forty-five minutes after exposure, compared to the unexposed control. Lastly, the samples that were subsequently exposed to 390-nm light decreased in bioluminescence forty-five minutes post exposure in comparison to those that were unexposed and those that were exposed only to 342 nm. As an additional control, the G69C mutant was exposed alongside Were-1 and showed no significant difference when G69C excited with 342 nm light, or 342 nm and 390 nm (Fig. 3f). These results strongly suggest that Were-1 is a photoriboswitch that can reversibly regulate protein expression *in vivo*.

Taken together, we show that novel riboswitches, regulated by synthetic ligands, can be evolved from random libraries fused to expression platforms. In the case of Were-1, binding of the target ligand stabilizes the RNA in a conformation that impedes the translation of a downstream ORF.

This approach will likely also yield transcription-regulating riboswitches, and further molecular engineering will allow regulation of a wide range of cellular events in both *cis* and *trans*. We expect that Were-1 and similar photoriboswitches will allow reversible photoregulation of a variety of RNA-centered cellular events with a very high spatiotemporal resolution in bacteria and multicellular organisms alike.

## MATERIALS AND METHODS

### Reagents and equipment

Unless otherwise stated, all reagents were purchased from Sigma-Aldrich. (E)-6’-(2-aminoethoxy)-2,2’,3,3’ tetrahydro-[1,1’-biindenylidene]-6-ol (amino-*t*SS) was synthesized and prepared as described below. Commercially available reagents were used without further purification. Absorbance spectra were recorded with a Thermo Scientific NanoDrop 1000 spectrophotometer. Fluorescence excitation and emission spectra were measured with a Varian Cary Eclipse fluorescence spectrometer, unless otherwise specified. Bioluminescence was measured using an Andor 866 EMCDD camera, BioTek Synergy H1 plate reader, or IVIS Lumina II.

### Synthesis of (E)-6’-(2-aminoethoxy)-2,2’,3,3’ tetrahydro-[1,1’-biindenylidene]-6-ol

All starting reagents were commercially available, and of analytical purity, and were used without further treatment. Solvents were dried according to standard methods. ^1^H and spectra were recorded on Varian UNITY INOVA-300 and Bruker Avance-600 instruments. Chemical shifts (*δ*) are reported in ppm relative to residual solvent peak (DMSO: *δ*_H_ = 2.50 ppm) as internal standard. Accurate mass measurements (HRMS) were obtained by ESI on an Agilent 6530 Q-TOF MS spectrometer. Analytical TLC was performed using a precoated silica gel 60 Å F_254_ plates (0.2 mm thickness) visualized with UV at 254 nm. Preparative column chromatography was carried out using silica gel 60 Å (particle size 0.063–0.200 mm). Purifications by HPLC were performed under the following conditions: Agilent ZORBAX SB-C18 column (5 µL, 9.4×150 mm); UV/Vis detection at λ_obs_ = 254 nm; flow rate 4 mL/min; gradient elution method H_2_O (0.1 % TFA) – CH_3_CN (0.1 % TFA) from 95:5 to 0:100 in 20 min. Purity of compounds was confirmed using Agilent eclipse plus C18 column (3.5 µL, 4.6×100 mm); UV/Vis detection at λ_obs_ = 254 nm; flow rate 0.5 mL/min; gradient elution method H_2_O (0.1 % TFA) – CH_3_CN (0.1 % TFA) from 95:5 to 0:100 in 20 min.

#### Synthesis details

**Scheme S1.**
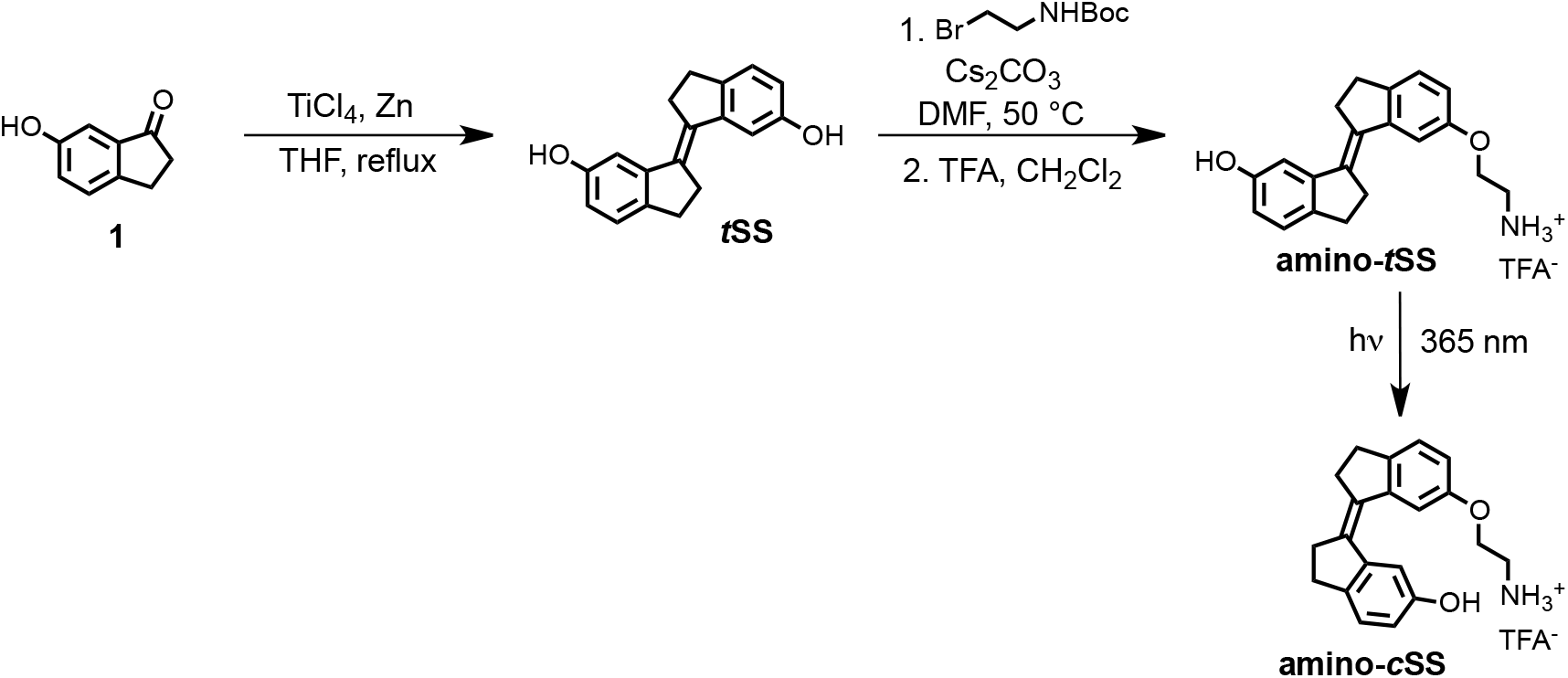
Synthesis of stiff stilbene derivatives ***t*SS, amino-*t*SS,** and **amino-*c*SS**.

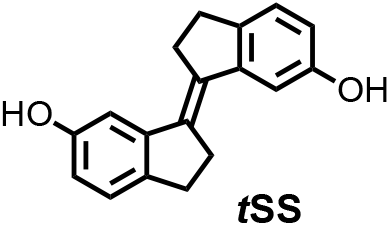

### (*E*)-2,2’,3,3’-tetrahydro-[1,1’-biindenylidene]-6,6’-diol (*t*SS)

To a stirred suspension of zinc powder (3.20 g, 48.92 mmol) in dry THF (50 ml), TiCl_4_ (4.67 g, 24.58 mmol) was added over 10 minutes at 0 °C. The resulting slurry was heated at reflux for 1.5 h. Then a THF solution (50 ml) of indanone **1** (600 mg, 4.05 mmol) was added over 3 h period by syringe pump to the refluxing mixture. The reflux was continued for 30 minutes after the addition was complete. After cooling to room temperature, the reaction mixture was poured into a saturated solution of NH_4_Cl and extracted with CH_2_Cl_2_. The organic solutions were dried over MgSO_4_ and concentrated by rotary evaporation under reduced pressure. The crude product was purified by column chromatography using silica gel (hexane/iPrOH = 10:0.5) to afford ***t*SS** in 51 % yield (273 mg) as a white solid. 1H NMR (400 MHz, DMSO-*d_6_*): *δ* 2.95 (m, 4H), 3.02 (m, 4H), 6.63 (dd, *J* = 8.1, 2.2 Hz, 2H), 7.02 (d, *J* = 2.3 Hz, 2H), 7.11 (d, *J* = 8.0 Hz, 2H), 9.17 (s, 2H). ^13^C NMR (100 MHz, DMSO-*d_6_*): *δ* 29.6, 31.9, 111.0, 114.5, 125.2, 135.0, 137.0, 143.7, 156.1. HRMS (ESI): *m/z* [M]^+^ calculated for C_18_H_16_O_2_ 264.1145; found: 264.1141.

### *E*)-6’-(2-aminoethoxy)-2,2’,3,3’-tetrahydro-[1,1’-biindenylidene]-6-ol (amino-*t*SS)

A mixture of ***t*SS** (125 mg, 0.47 mmol), 2-(Boc-amino)ethyl bromide (106 mg, 0.47 mmol),

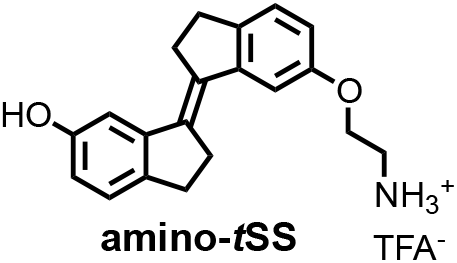

Cs_2_CO_3_ (772 mg, 2.37 mmol) and *n*Bu_4_NBr (6 mg, 0.02 mmol) in DMF (15 ml) was heated at 50 °C for 6 h. After cooling to room temperature, CH_2_Cl_2_ was added and the mixture was washed with a saturated solution of NH_4_Cl, water, brine and dried over MgSO_4_. The organic solutions were concentrated by rotary evaporation under reduced pressure. The crude product was dissolved in a mixture of CH_2_Cl_2_/TFA (16 ml; 3:1) and stirred at room temperature for 30 minutes. The reaction mixture was concentrated by rotary evaporation under reduced pressure and purified by HPLC (gradient elution method H_2_O (0.1 % TFA) – CH_3_CN (0.1 % TFA) from 95:5 to 0:100) to afford **amino-*t*SS** (as a TFA salt) in 42 % yield (84 mg) as an off-white solid. 1H NMR (400 MHz, DMSO-*d_6_*): *δ* 2.96 (m, 4H), 3.01 (m, 4H), 3.22 (t, *J* = 5.1 Hz, 2H), 4.17 (t, *J* = 5.1 Hz, 2H), 6.66 (dd, *J* = 8.1, 2.2 Hz, 1H), 6.87 (dd, *J* = 8.2, 2.3 Hz, 1H), 7.03 (d, *J* = 2.2 Hz, 1H), 7.13 (d, *J* = 8.1 Hz, 1H), 7.16 (d, *J* = 2.4 Hz, 1H), 7.27 (d, *J* = 8.3 Hz, 1H), 8.10 (s, 3H), 9.28 (s, 1H). ^13^C NMR (100 MHz, DMSO-*d_6_*): *δ* 29.58, 29.64, 31.8, 31.9, 38.5, 54.7, 110.8, 111.1, 113.6, 114.8, 125.3, 125.4, 134.5, 135.9, 137.1, 139.6, 143.6, 144.0, 156.2, 156.9. HRMS (ESI): *m/z* [M+H]^+^ calculated for C_20_H_21_NO_2_ 308.1645; found: 308.1651.

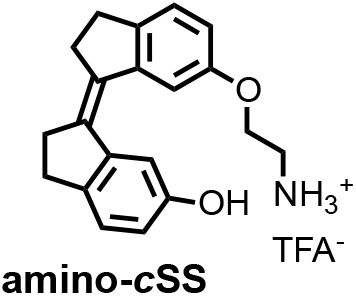

### *(Z*)-6’-(2-aminoethoxy)-2,2’,3,3’-tetrahydro-[1,1’-biindenylidene]-6-ol ((Z)-1; amino-*c*SS)

A solution of **amino-*t*SS** (15 mg, 35.6 µmol) in DMSO (1 ml) in NMR cuvette was irradiated with handheld UV lamp (8 W) for 15 min. The resulting mixture of **amino-*t*SS** and **amino-*c*SS** was purified by HPLC (gradient elution method H_2_O (0.1 % TFA) – CH_3_CN (0.1 % TFA) from 95:5 to 0:100) to afford **amino-*c*SS** (as a TFA salt) in 20 % yield (2 mg) as an off-white solid. ^1^H NMR (400 MHz, DMSO-*d_6_*): *δ* 2.75 (m, 4H), 2.85 (m, 4H), 3.22 (t, *J* = 5.1 Hz, 2H), 4.13 (t, *J* = 5.1 Hz, 2H), 6.63 (dd, *J* = 8.2, 2.2 Hz, 1H), 6.84 (dd, *J* = 8.2, 2.2 Hz, 1H), 7.12 (d, *J* = 8.2 Hz, 1H), 7.25 (d, *J* = 8.3 Hz, 1H), 7.40 (d, *J* = 2.0 Hz, 1H), 7.53 (d, *J* = 2.0 Hz, 1H), 8.02 (s, 3H), 9.25 (s, 1H). ^13^C NMR (100 MHz, DMSO-*d_6_*): *δ* 29.2, 29.3, 34.9, 35.0, 38.5, 64.4, 108.9, 109.4, 114.5, 115.2, 125.8, 125.9, 134.4, 135.7, 138.6, 140.79, 140.84, 141.1, 155.46, 156.1. HRMS (ESI): *m/z* [M+H]^+^ calcd for C_20_H_21_NO_2_ 308.1645; found: 308.1644.

### *In vitro* RNA transcription

RNA was transcribed at 37 °C for one hour in a 50 µL volume containing 40 mM tris-HCl, 6 mM dithiothreitol (DTT), 2 mM spermidine, 1.25 mM each rNTP, 8 mM MgCl_2_, 1 unit of T7 RNA polymerase, and 5 pmol of DNA template. The transcripts were purified by 10 % PAGE under denaturing conditions (7 M urea). RNA was eluted from the gel into 300 µL of 300 mM KCl and precipitated by adding 700 µL of 95 % ethanol at –20 °C.

### *In vitro* selection of amino-*t*SS aptamers

An RNA pool derived from a *B. subtilis mswA* SAM-1 riboswitch, located in the 5≠ untranslated region of the *metI* (cystathionine gamma-synthase, also denoted as *yjcI*) gene ^24, 33^ was designed by replacing the riboswitch ligand-binding domain with a random region of 45 nucleotides. The anti-terminator stem and upstream half of the transcriptional terminator sequence were partially randomized at a 15 % level, and the loop of the terminator stem was fully randomized. The remaining part of the riboswitch, including the downstream half of the transcriptional terminator stem, containing a ribosome binding site (RBS) that binds the 3’ end of *B. subtilis* 16S rRNA (3’-UUUCCUCCACUAG-5’) ^34^ and an alternative UUG start codon, was retained (Supplementary Fig. 1). The pool was synthesized by Yale School of Medicine’s Keck Oligonucleotide Synthesis facility as a single template strand that was then purified by 10 % PAGE and converted into dsDNA by a primer-extension reaction using a primer corresponding to the T7 RNA polymerase promoter. The pool was transcribed at an estimated sequence diversity of 10^15^.

From that pool, RNAs were selected to bind amino-*t*SS, as follows. PAGE-purified ^32^P-labeled RNA transcripts of the pool were precipitated, dried, and resuspended in a solution containing 140 mM KCl, 10 mM NaCl, 10 mM tris-chloride, pH 7.5, and 5 mM MgCl_2_ (binding buffer). The RNA mixture was heated to 70 °C for three minutes and loaded onto agarose beads for a counter-selection step. Binders were discarded and the flow-through was incubated on agarose beads linked to amino-*t*SS. The beads were shaken for five minutes at room temperature, and the unbound RNA was collected. Amino-*t*SS beads were then washed with binding buffer for five minutes at room temperature. This washing step was repeated six times. Potential aptamers were then eluted twice with denaturing buffer, consisting of 7 M urea and 5 mM ethylenediaminetetraacetic acid (EDTA) in 45 mM tris, 45 mM borate buffer, pH 8, and heated at 95 °C for five minutes. Each fraction was analyzed for radioactivity using a liquid scintillation counter. Elutions were pooled, precipitated, dried, and resuspended in water for reverse transcription. The pool was reverse transcribed, and the cDNA was amplified by PCR and used for the next round of selection.

### Screening of potential amino-*t*SS binders

After six rounds of *in vitro* selection, the selected pool was cloned into a TOPO TA plasmid (Invitrogen) and transformed into DH5α *E. coli* cells. Cells were plated on agar containing kanamycin and incubated overnight at 37 °C. Individual colonies were picked from the master plate and inoculated overnight in Luria Broth containing kanamycin. Plasmids were extracted and purified using a Miniprep Kit (QIAGEN), and sequenced (GENEWIZ). Individual clones were PCR-amplified using the library-specific primers and transcribed to test their optical activity in the presence and absence of amino-*t*SS. *t*SS emission spectra were collected using an excitation at 355 nm. Were-1 showed the highest increase in amino-*t*SS fluorescence at 430 nm and was chosen for further analysis.

#### Structure probing of Were-1

#### T1 nuclease probing

Were-1 RNA was dephosphorylated in a solution of the reaction buffer (50 mM potassium acetate, 20 mM Tris-acetate, 10 mM magnesium acetate, 100 µg/ml BSA, pH 7.9), 1 µg of purified RNA, and 1.5 unit of Shrimp Alkaline Phosphatase (NEB). The reaction was incubated at 37 °C for 30 minutes, and heat-inactivated at 65 °C for 5 minutes.

5′–labeled RNA (8000 cpm) was prepared in reaction buffer (70 mM Tris-HCl, pH 7.6, 10 mM MgCl_2_, 5 mM DTT) using 1 µg of 5′–dephosphorylated RNA, 2 µCi [γ-^32^P] ATP (Perkin Elmer), and 15 units of T4 PNK enzyme (NEB). The reaction was incubated at 37 °C for two hours and PAGE–purified.

The 5′–labeled Were-1 RNA was added into binding buffer and the indicated concentrations of amino-*t*SS or controls (no ligand, amino-*c*SS, tS, tDHS, and SAM), and were incubated at 55 °C for 5 minutes and subsequently cooled at room temperature for 5 minutes. Next, T1 nuclease was added (0.05 units; Thermo Fisher Scientific) and samples were incubated at 37 °C for 15 minutes. All conditions were then quenched with a mixture of 7 M urea and 10 mM EDTA. Afterwards, the RNA was added to an equal volume of phenol:chloroform:isoamyl alcohol (25:24:1) and vortexed. Samples were centrifuged for 3 minutes at 8,000 RPM, and the aqueous phase was collected and transferred to a new tube. Samples were fractionated on a 10% PAGE gel and exposed to a phosphor image screen (GE Healthcare) for a minimum of 24 hours. The screen was scanned on a GE Typhoon phosphor imager.

A guanosine-specific sequencing lane was resolved in parallel to all samples using 5′–labeled or 3′–labeled RNA (8000 cpm), as specified, in T1 digestion buffer (250 mM sodium citrate, pH 7) and 0.5 units T1. Reactions were incubated at 55 °C for 5 minutes and quenched with a solution containing 7 M urea and 10 mM EDTA. RNA was extracted using phenol-chloroform, as noted above. Partial alkaline hydrolysis was also resolved in parallel by adding 5′–labeled or 3′–labeled RNA (8000 cpm), as specified, into a hydrolysis buffer (50 mM NaHCO_3_, 1 mM EDTA, pH 10). Reactions were incubated at 95 °C for 10 minutes and quenched in a solution containing 7 M Urea and 10 mM EDTA. RNA was extracted using phenol-chloroform.

### SHAPE

A selective 2’-hydroxyl acylation and primer extension (SHAPE) reaction, as described 35, was carried out on Were-1 in the presence of increasing amino-*t*SS concentrations and 30 µM controls (amino-*c*SS, tDHS, tS, and SAM).

#### S1 nuclease probing

Reactions were prepared by adding 3′–labeled Were-1 RNA (8000 cpm) into S1 nuclease buffer (40 mM sodium acetate, pH 4.5, 300 mM NaCl, and 2 mM ZnSO_4_), and the indicated concentrations of amino-*t*SS, and were incubated at 55 °C for 5 minutes and subsequently cooled at room temperature for 5 minutes. Next, S1 nuclease was added (0.2 units; Thermo Fisher Scientific) and samples were incubated at 37 °C for 10 minutes and quenched in a solution of 7 M Urea and 10 mM EDTA. Samples were extracted using phenol-chloroform and resolved on a denaturing 10 % PAGE gel. The gel was then exposed to a phosphor image screen and scanned on a GE Typhoon phosphor imager. The sequences in the degradation pattern were assigned by running T1 digestion and partial alkaline hydrolysis in parallel lanes, as noted above.

#### Terbium (III) footprinting

Reactions were prepared by adding 5′–labeled Were-1 RNA (8000 cpm) into the binding buffer, and the indicated concentrations of amino-*t*SS or controls (no ligand, amino-*c*SS, tS, tDHS, and SAM), and were incubated at 55 °C for 5 minutes and subsequently cooled at room temperature for 5 minutes. Terbium (III) chloride was added to a final concentration of 10 mM and samples were incubated at 37 °C for 30 minutes and then quenched with a solution of 7 M urea and 10 mM diethylenetriaminepentaacetic acid (DTPA). Were-1 RNA was extracted using phenol-chloroform, as noted above, and samples were fractionated on a denaturing 10 % PAGE gel. The gel was exposed to a phosphor image screen (GE Healthcare) and scanned on a GE Typhoon phosphor imager. The sequences in the degradation pattern were assigned by running TI digestion and partial alkaline hydrolysis in parallel lanes, as noted above.

#### In–line probing

Reactions were prepared by adding 3′–labeled RNA (8000 cpm) into the binding buffer, pH 8.5, and the indicated concentrations of controls (no ligand, cSS, tS, tDHS, and SAM). Samples were initially incubated at 55 °C for 5 minutes and then cooled at room temperature for 5 minutes, and then incubated at 37 °C for 20 hours. All conditions were quenched in a solution of 7 M Urea and 10 mM EDTA. Were-1 RNA was extracted using phenol-chloroform, as noted above, and run on a denaturing 10 % PAGE gel. The gel was exposed to a phosphor image screen (GE Healthcare), and scanned on a GE Typhoon phosphor imager. The sequences in the degradation pattern were assigned by running T1 digestion and alkaline hydrolysis in parallel lanes, as noted above.

All gels were analyzed in ImageJ. Structure predictions of Were-1 in the absence of amino-*t*SS were performed using RNAfold of the Vienna RNA package ^36^ (Fig. 1c).

### *In vitro* strand displacement reaction

A dsDNA reporter was designed to contain a toehold that complements the Shine-Dalgarno sequence of the riboswitch, in which the longer (toehold) strand (Rep F) contained the 3’ toehold sequence, a reverse complement of the Shine-Dalgarno sequence, as well as a 5’ fluorescein. The shorter strand (Rep Q) contained a 3’ Iowa black quencher (Supplementary Table 1). A solution of 2:1 Rep Q:Rep F oligos in binding buffer was incubated at 95 °C for 1 minute, followed by 25 °C for 5 minutes, to anneal the strands and form the dsDNA reporter construct.

In a Falcon 384-well Optilux Flat Bottom plate, strand displacement was initiated by adding 100 nM of purified Were-1 RNA to 50 nM of toehold-fluorophore reporter. Amino-*t*SS was quickly added to some samples to test for ligand-dependent displacement. Fluorescence emission was recorded in a BioTek Synergy plate reader over a 45-minute period under continuous illumination using the following parameters: excitation wavelength, 485 nm; emission wavelength, 520 nm.

### *In vitro* co–transcriptional toehold-binding kinetics of Were-1

*In vitro* transcription was performed similarly to the above–described RNA transcription assay with the following modifications: 3 pmol template DNA and 50 nM toehold-fluorophore reporter were used. A 30 µL transcription reaction was initiated by the addition of 4 mM rNTP mix (containing 1mM of each rNTP) and fluorescence emission of the toehold-fluorophore reporter was recorded in a Varian Cary Eclipse fluorimeter under continuous illumination at 37 °C using the following parameters: excitation wavelength, 485 nm; emission wavelength, 520 nm; increment of data point collection, 0.01 s; slit widths, 10 nm. These conditions were used for the entire experiment unless stated otherwise. After an initial fluorescence increase, corresponding to the initial burst of transcription, amino-*t*SS was rapidly added to the solution and fluorescence emission was recorded for 200 s. To switch amino-*t*SS to the *cis* isoform (amino-*c*SS), the solution was excited at 342 nm (slit width, 2.5 nm; Φ_q_ = 6.8*10^-5^ W/cm^2^) for 60 s. Fluorescence emission of the toehold-fluorophore reporter was again recorded for 200 s. To switch the *cis* isoform back to the *trans* state, the solution was excited at 372 nm (slit width, 2.5 nm; Φ_q_ = 10*10^-5^ W/cm^2^) for 60 s. Again, fluorescence emission of the toehold-fluorophore reporter was recorded for 200 s. This process was repeated two to three more times until fluorescence plateaued.

### IC_50_ measurements

A dose-response of the Were-1 riboswitch to the target ligand (amino-*t*SS) was assessed by measuring fluorescence as a function of ligand concentration in the presence of a toehold-fluorophore reporter construct (50 nM). Fluorescence emission was recorded under continuous illumination at 37 °C using the following parameters: excitation wavelength, 485 nm; emission wavelength, 520 nm; increment of data point collection, 0.01 s; slit widths, 10 nm. The apparent rate constants were measured and plotted against the amino-*t*SS concentrations (or other ligands, as specified). The data were normalized to the no-amino-*t*SS control. The IC_50_ was extracted from fitting a curve to the graph using the equation:

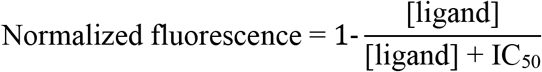

### *In vitro* co–transcriptional magnesium dependence of Were-1 toehold-binding

Using the same conditions as above, the fluorescence response of toehold-binding to the Were-1 riboswitch in the presence of 8.4 µM amino-*t*SS under various Mg^2+^ concentrations was measured. Fluorescence emission was recorded under continuous illumination at 37 °C on a BioTek Synergy H1 plate reader.

### Cloning the Were-1 riboswitch for expression in *E. coli* cells

Were-1 DNA was cloned into the pBV-Luc (Addgene) vector in order to obtain a fused riboswitch-firefly luciferase (Fluc) reporter construct. The PCR primers were designed to add a 5’ *EcoRI* site to the template Were-1 DNA upstream of the T7 promoter and a 3’ overhang containing 35 nucleotides of the Fluc gene directly downstream of its start codon to replace the Fluc start codon sequence. Both the PCR product and plasmid were digested by *EcoRI HF* and *KasI* (New England BioLabs) and purified. The purified construct was then inserted at the 5’ end of the Fluc coding sequence with T4 DNA ligase (New England BioLabs). The resulting vector was termed Were-1-Fluc (Table 1).

Were-1-Fluc was transformed into DH5α *E. coli* cells and grown overnight on agar plates containing ampicillin at 37 °C. Ten colonies were picked from a master plate and individual clones were inoculated overnight in Luria Broth containing ampicillin. Plasmids were purified using a Miniprep Kit (QIAGEN) and individually sequenced (GENEWIZ). Correct constructs were transcribed *in vitro* and fractionated on an agarose gel to confirm sequencing results by measuring the size of the fused construct. Using the same procedure as above, one clone was analyzed in an *in vitro* co-transcriptional toehold-binding experiment to test whether the new fused construct was able to function similarly to the stand-alone riboswitch.

### *In vitro* transcription and translation kinetics

The PURExpress *in vitro* protein synthesis kit (New England BioLabs) was used to transcribe and translate Were-1-Fluc. Experiments were performed similarly to the kit assay conditions with the following modifications: 200 ng/µL DNA, 100 µM D-luciferin, and 2 mM MgCl_2_. Amino-*t*SS (or other ligands, as specified) was added in conditions when specified. A control plasmid, pET-Luc2, was also tested in the presence and absence of 11 µM amino-*t*SS. All luminescence data were acquired using an ANDOR camera (EMCCD) at 25 °C and analyzed using Solis software, and images were further processed and analyzed using ImageJ.

To test whether Were-1 could regulate luciferase protein expression, samples were prepared under identical conditions and luminescence was measured for approximately 40 mins. Samples were then excited at 342 nm (Φ_q_ = 1.4*10^-2^ W/cm^2^) for 1 s, and luminescence was recorded for approximately 30 mins. Samples were excited at 390 nm (Φ_q_ = 5.5*10^-2^ W/cm^2^) for 1 s to switch Were-1 back to the bound ‘off’ state, and again, luminescence was measured for approximately 30 mins.

### IC_50_ measurements

A dose-response of the Were-1 riboswitch to amino-*t*SS was assessed by measuring luminescence as a function of increasing target concentration. All data were acquired using an ANDOR camera and analyzed with Solis software, and images were further processed and analyzed using ImageJ, as described above.

### *In vivo* translation kinetics

Were-1-Fluc was transformed into BL21(DE3) *E. coli* cells and grown overnight in Luria Broth containing ampicillin (OD_600_ = 0.26). 1 mM IPTG was added to each well (containing 45 µL culture) to induce T7 RNA polymerase-driven expression and 100 µM D-luciferin to provide a substrate for Fluc. Amino-*t*SS or amino-*c*SS was also added where specified. Bioluminescence was recorded every five mins for one hour at 37 °C using a BioTek Synergy H1 plate reader.

To test whether the Were-1 riboswitch regulates the production of Fluc, samples were prepared under the same conditions. In the presence of amino-*t*SS, bioluminescence was measured on a BioTek Synergy H1 plate reader for approximately 15 mins before samples were excited at 342 nm for 1s in order to isomerize amino-*t*SS to amino-*c*SS. Luminescence was recorded again for approximately 20 mins. Similar experiments were used regarding amino-*c*SS, with the exception of using 390 nm exposure for 0.5 ms in order to isomerize amino-*c*SS to amino-*t*SS, unless further specified.

### *In vivo* light exposure analysis

To determine the dependence of the amino-*t*SS exposure on the Were-1-Fluc expression, samples were prepared as described above and loaded into two black-bottom 96-well plates. One plate was used as a control and the other was exposed to 342 nm light (Φ_q_ = 1.4*10^-2^ W/cm^2^) for their specified time using a Nikon FM-10 camera shutter in order to isomerize amino-*t*SS to amino-*c*SS. The same procedure was performed in the presence of amino-*c*SS, except 390 nm light was used (Φ_q_ = 5.5*10^-2^ W/cm^2^) for their specified time to isomerize amino-*c*SS to amino-*t*SS. Bioluminescence was measured on an IVIS Lumina II imaging system 1 hr after light exposure.

To test whether the Were-1 riboswitch regulates the production of Fluc multiple times, samples were prepared as described above and loaded in a black-bottom 96-well plate. The top half of the plate was used as a control, containing the unresponsive G69C mutant, and the bottom half contained Were-1. All samples contained 10 *µ*M amino-*t*SS and had their bioluminescence was measured 25 minutes after induction on a BioTek Synergy H1 plate reader. Next, the first group of wells remained unexposed to light and the middle and far right samples were exposed to 342 nm (Φ_q_ = 1.4*10^-2^ W/cm^2^) for 1 ms. Measurements for all samples were taken 40 minutes after 342 nm exposure. Finally, the last group of wells (far right) were excited at 390 nm for 0.5 ms. Bioluminescence was measured again, for all samples, 40 minutes after exposure. The same experiment was repeated with the inactive mutant, G69C, as an additional control. Data were normalized to the unexposed samples, and OD_600_ values were obtained to confirm that there was no cell death from UV damage.

### IC_50_ measurements

A dose-response of the Were-1 riboswitch to the target metabolite (amino-*t*SS) was assessed by measuring bioluminescence inhibition as a function of increasing target concentration in BL21(DE3) *E. coli* cells. Bioluminescence was recorded under continuous conditions at 37 °C.

## Data Availability

The authors declare that all data are available in the manuscript or the supplementary materials.

## Acknowledgments

We thank the researchers who provided support for this study: Dalen Chan for synthesizing NAI, and Jennifer Prescher’s lab for their luminescence reagents and IVIS setup. This work was supported by grants from the National Science Foundation (1804220 to A.L.), the National Institutes of Health (5T32GM108561-04 to K.A. R. and 5R01GM094929 to A.L.), the John Templeton Foundation (through the Foundation for Applied Molecular Evolution to A.L.), and the Czech Science Foundation (GACR 17-25897Y to J.M.).

## Authors contributions

A. L. and J. M. designed the study. J. M. and M. M. A. designed the *in vitro* selection experiment. J. M. and K. S. synthesized the ligands with the help of A. R. C. J. M. characterized the ligand. M. M. A. carried out the *in vitro* selection. K. A. R. performed the *in vitro* and *in vivo* riboswitch characterization experiments. F. C. and K. A. R. designed the luciferase reporter system. L. F. M. P., K. A. R., and M. M. A. performed the structure probing of Were-1. A. L. and K. A. R. prepared figures and wrote the manuscript with input from all other authors.

