## Supplementary Figures and Table for "Regulation of mRNA translation by a photoriboswitch"

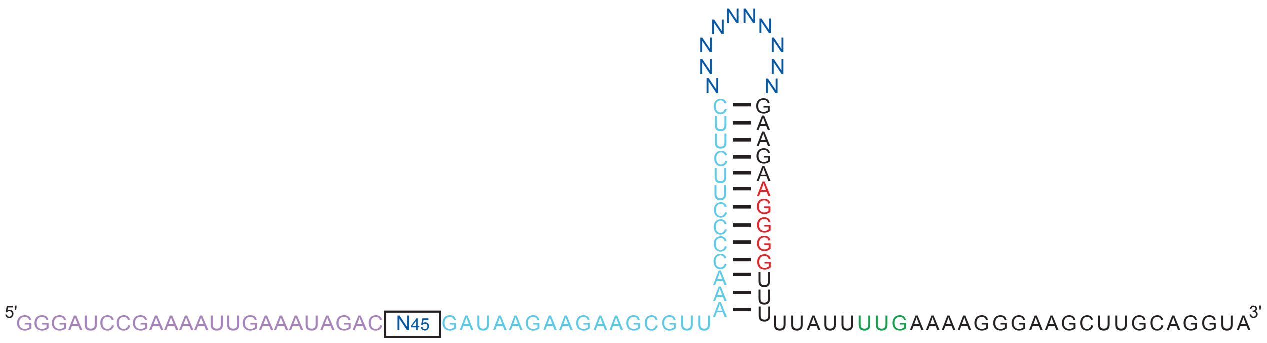


**Supplementary Figure 1 | *In vitro* selection pool design for a photoriboswitch.** An RNA pool was derived from *B. subtilis mswA* SAM-I riboswitch by replacing its ligand-binding domain with a 45-nucleotide random sequence (dark blue, boxed), partially randomizing (at a 15% level) its anti-terminator hairpin and the 5' half of the terminator hairpin (light blue), replacing the terminator-helix tetraloop sequence (UUAU) with a random decamer (dark blue), and retaining the 3' half of the terminator hairpin and its translation initiation sequences (red – Shine-Dalgarno; green – start codon). The pool's forward primer sequence (in RNA form) is shown in purple.

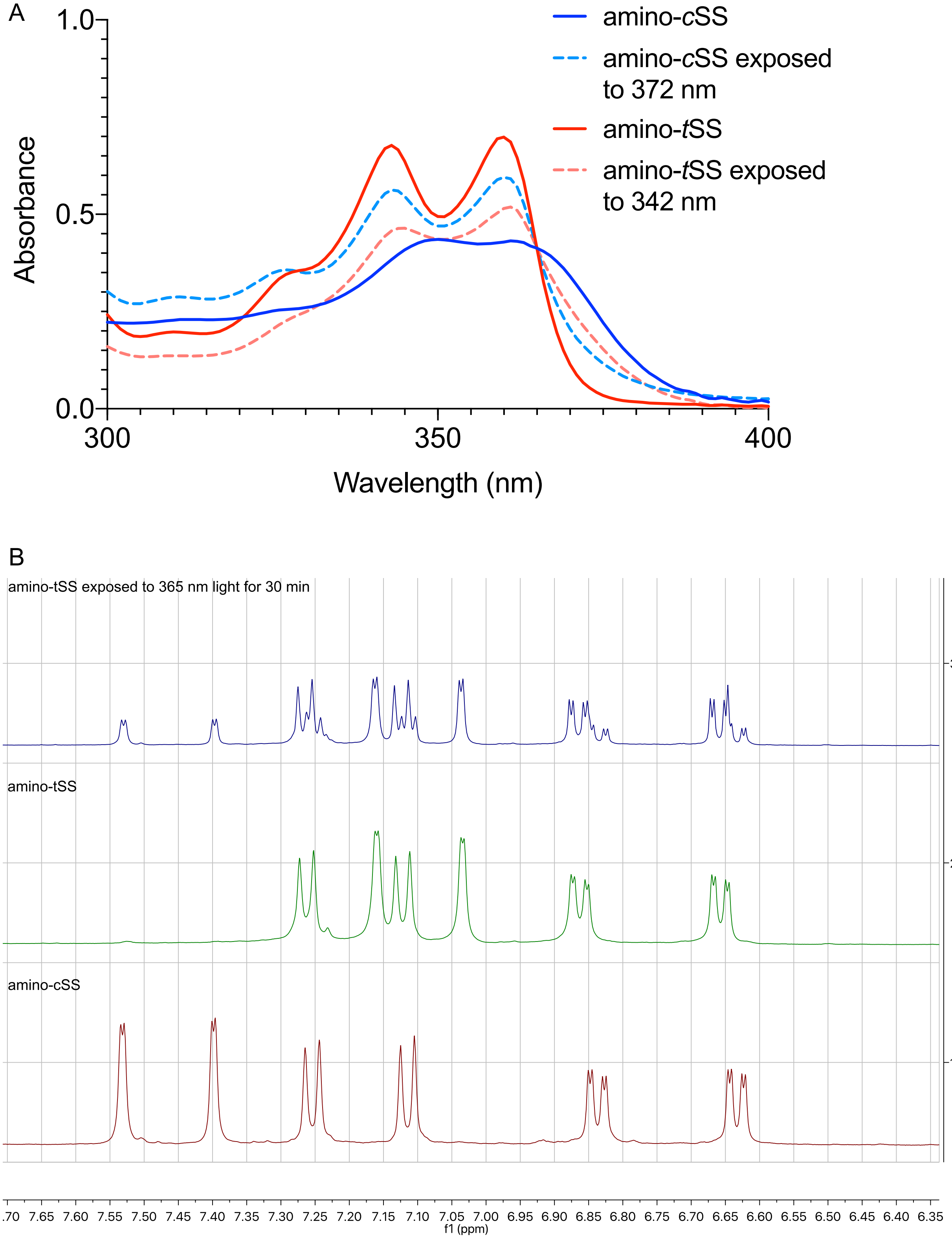
A
Absorbance
1.0
0.5
0.0
300
350
400
Wavelength (nm)
amino-cSS
amino-cSS exposed to 372 nm
amino-tSS
amino-tSS exposed to 342 nm
B
amino-tSS exposed to 365 nm light for 30 min
amino-tSS
amino-cSS
3
2
1
.70 7.65 7.60 7.55 7.50 7.45 7.40 7.35 7.30 7.25 7.20 7.15 7.10 7.05 7.00 6.95 6.90 6.85 6.80 6.75 6.70 6.65 6.60 6.55 6.50 6.45 6.40 6.35
f1 (ppm)

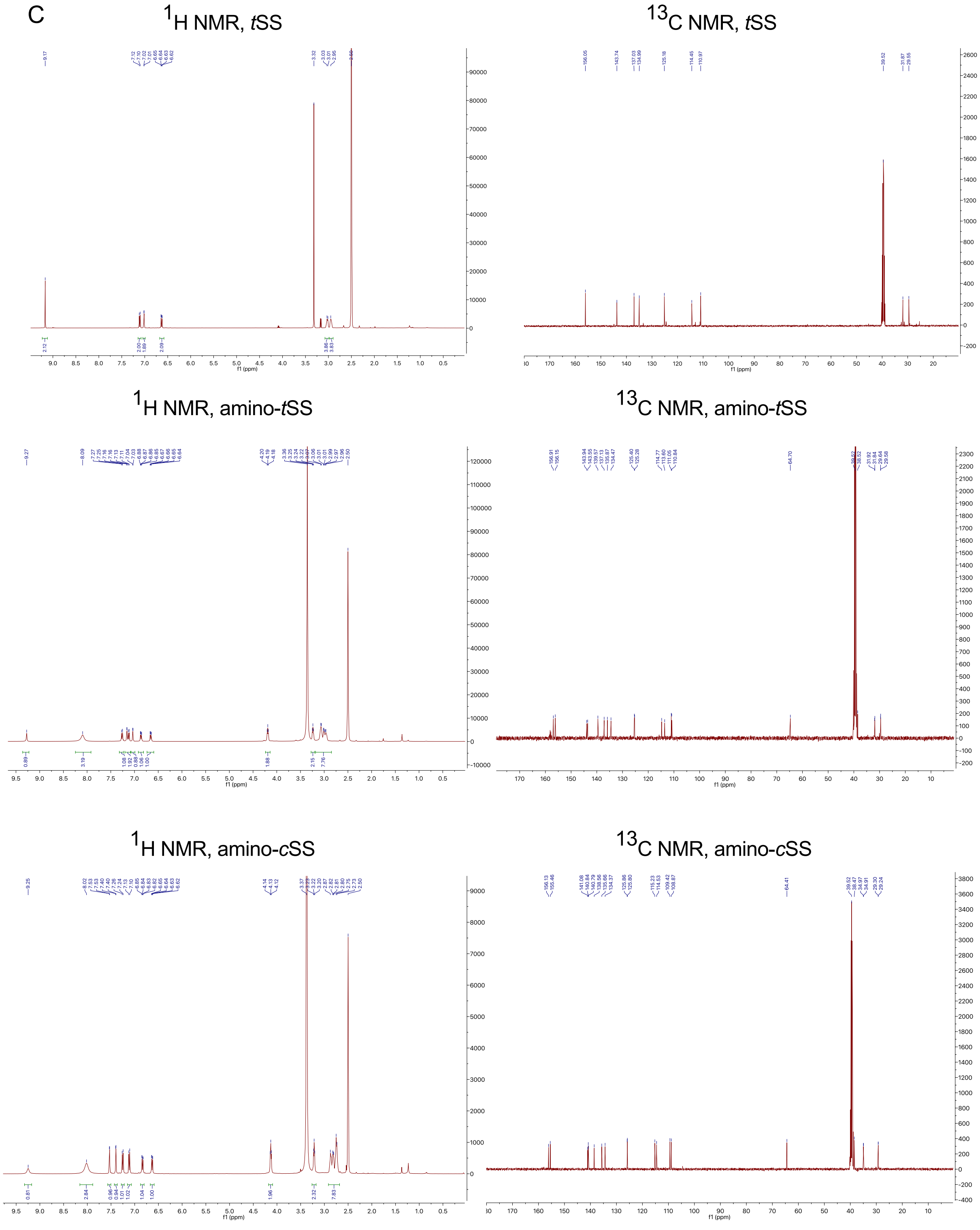
C
$^1$H NMR, *t*SS
$^{13}$C NMR, *t*SS
$^1$H NMR, amino-*t*SS
$^{13}$C NMR, amino-*t*SS
$^1$H NMR, amino-*c*SS
$^{13}$C NMR, amino-*c*SS
f1 (ppm)

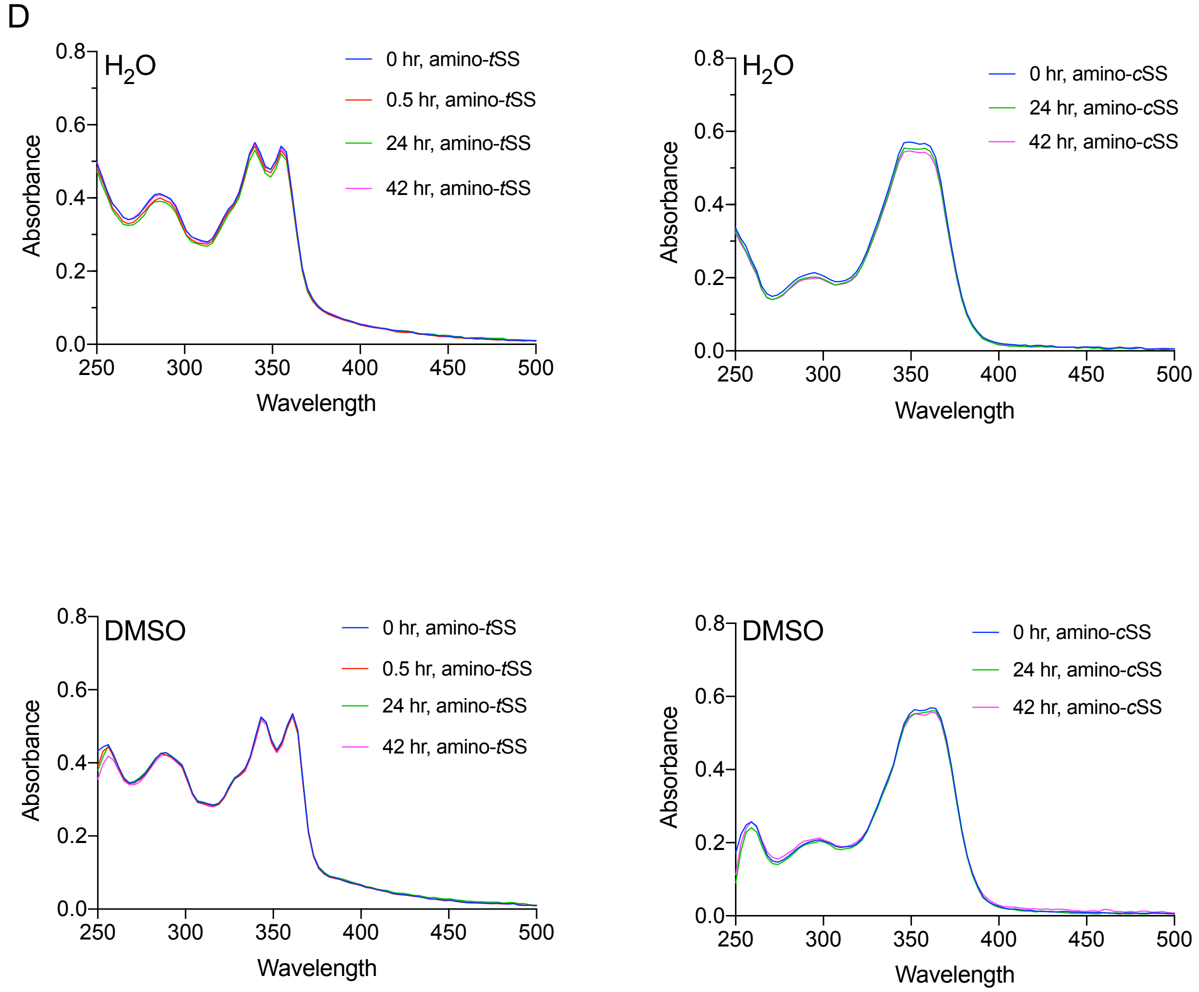


**Supplementary Figure 2 | E/Z isomerization of amino-*t*SS and amino-*c*SS. a**, UV-Vis spectra of amino-*t*SS (red) and amino-*t*SS after 3 h exposure to 342 nm (red, dashed), and amino-*c*SS (blue) and amino-*c*SS after 1.5 h exposure to 372 nm (blue, dashed). Photo isomerization was performed in 30 mM DMSO solutions and subsequently diluted 1000 times for UV-Vis spectra measurement. **b**, $^{1}$H NMR of amino-*t*SS, amino-*c*SS, and E/Z isomerization. At the photostationary state (amino-*t*SS exposed to 365 nm) the ratio of amino-*t*SS to amino-*c*SS is approximately 2:1 as determined by NMR. **c**, $^{1}$H NMR and $^{13}$C NMR spectra of *t*SS (top), amino-*t*SS (middle), and amino-*c*SS (bottom). **d,** UV-Vis spectra of amino-*t*SS and amino-*c*SS in

$H_2O$ (top) and DMSO (bottom). Both isoforms of the ligand are relatively stable in either solvent for the indicated times.

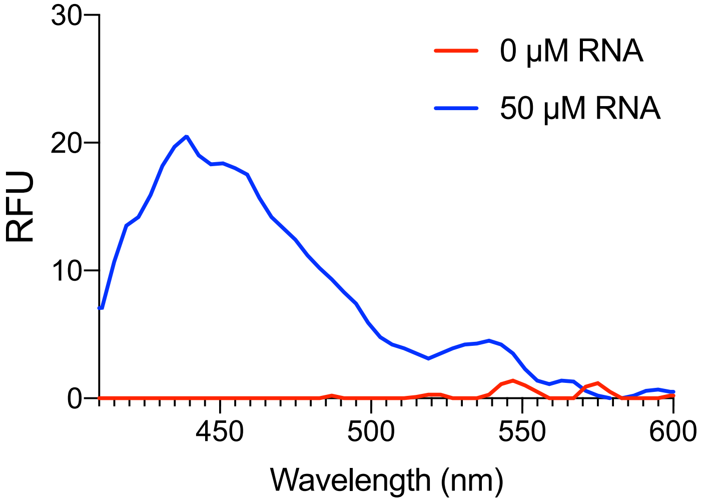


**Supplementary Figure 3 | Screening of Were-1 based on optical activity.** Fluorescence emission of 100 nM amino-*t*SS incubated with purified Were-1 RNA showing an increase in presence of Were-1 and suggesting RNA affinity for the target ligand. Emission spectra were collected using an excitation of 365/10 nm.

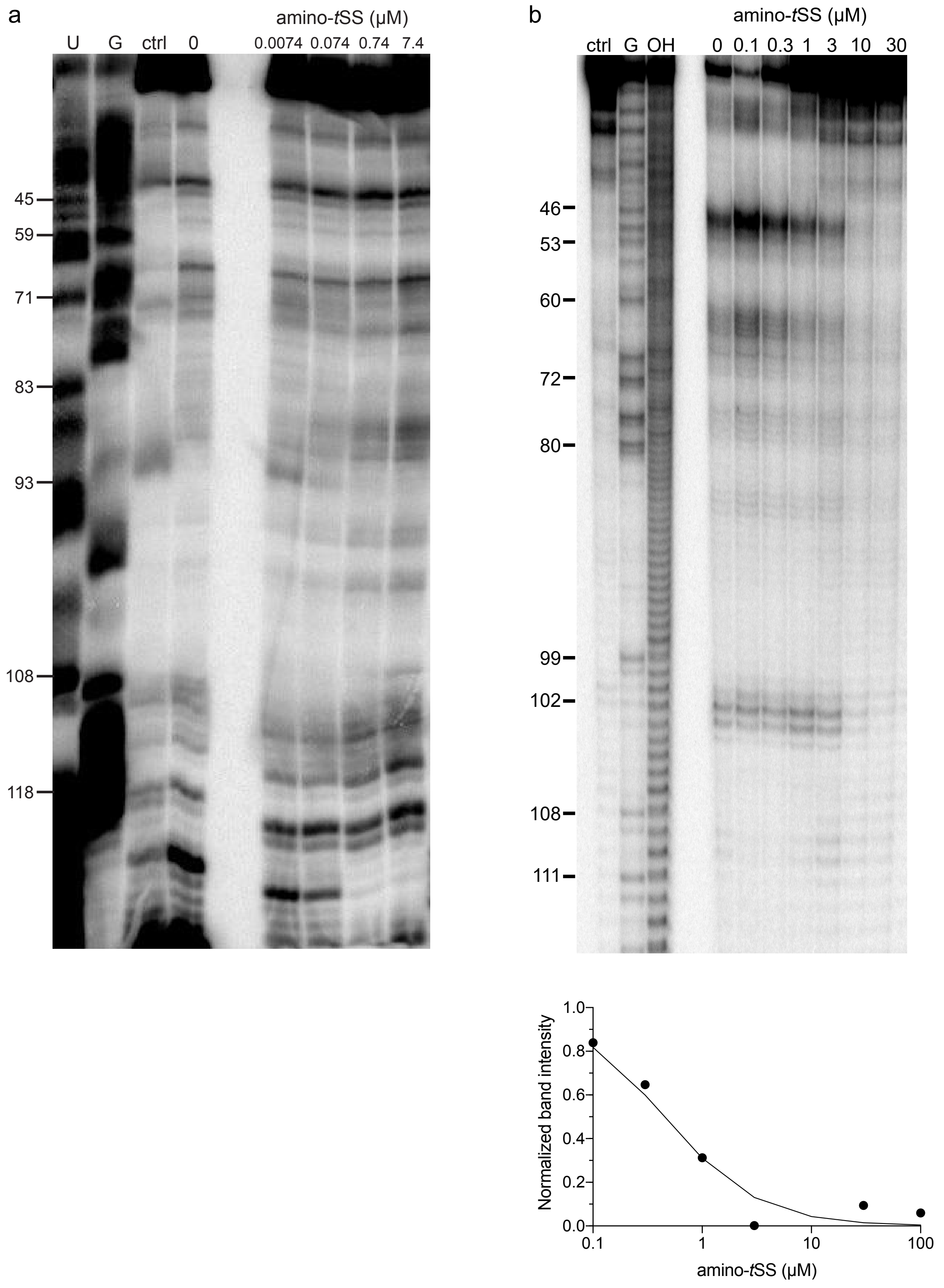
a
amino-tSS (µM)
U G ctrl 0 0.0074 0.074 0.74 7.4
45
59
71
83
93
108
118
b
amino-tSS (µM)
ctrl G OH 0 0.1 0.3 1 3 10 30
46
53
60
72
80
99
102
108
111
Normalized band intensity
1.0
0.8
0.6
0.4
0.2
0.0
0.1
1
10
100
amino-tSS (µM)

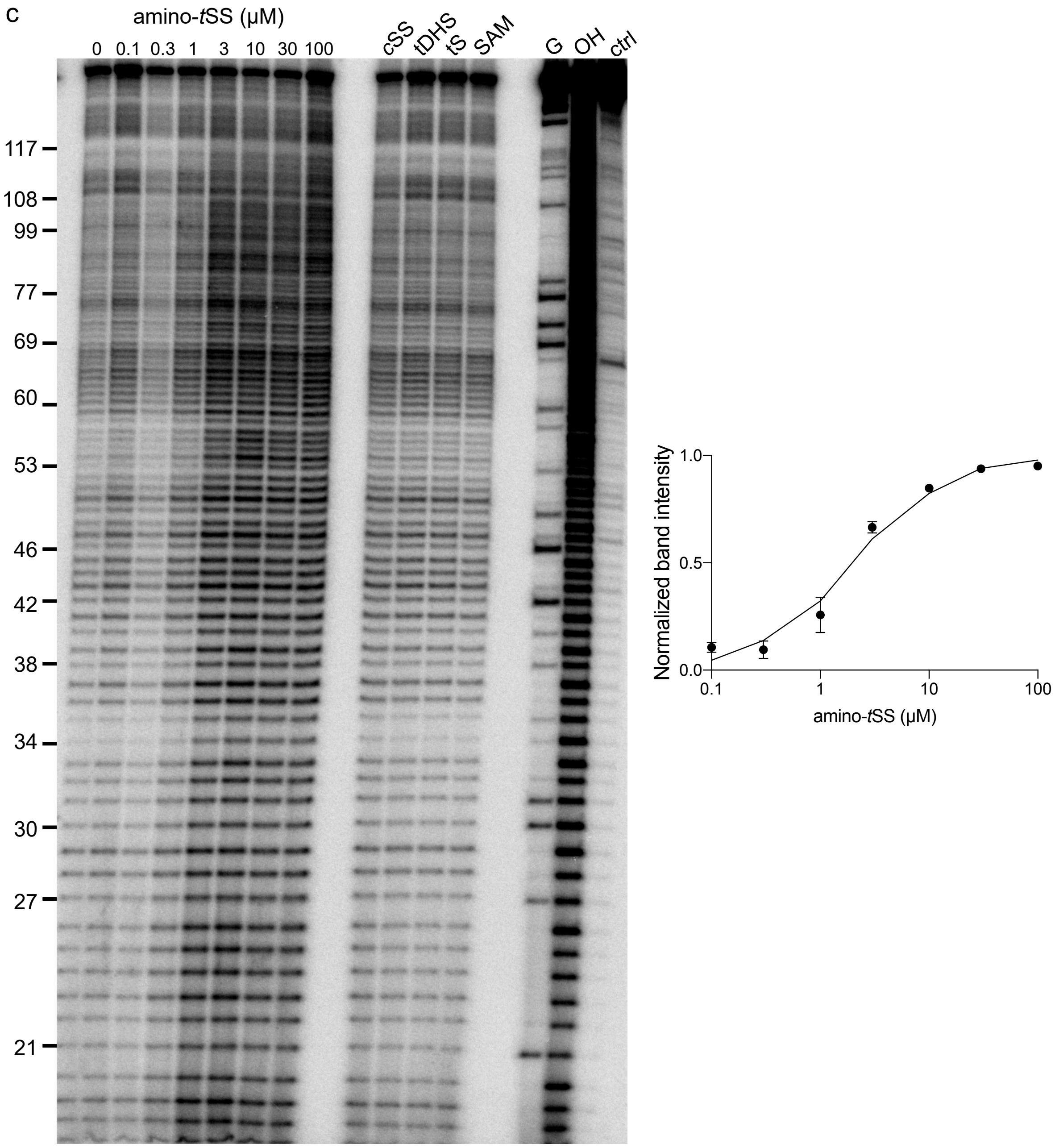
c
amino-tSS (μM)
0 0.1 0.3 1 3 10 30 100
cSS tDHS tS SAM
G OH ctrl
117
108
99
77
69
60
53
46
42
38
34
30
27
21
Normalized band intensity
1.0
0.5
0.0
0.1
1
10
100
amino-tSS (μM)

d

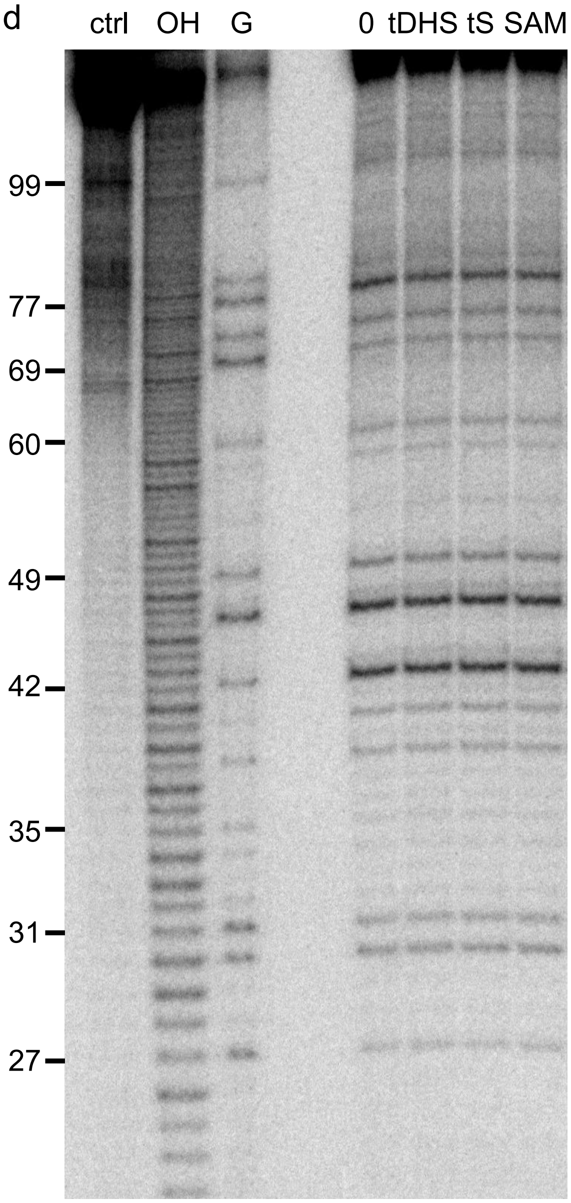
ctrl
OH
G
0
tDHS
tS
SAM
99
77
69
60
49
42
35
31
27

e

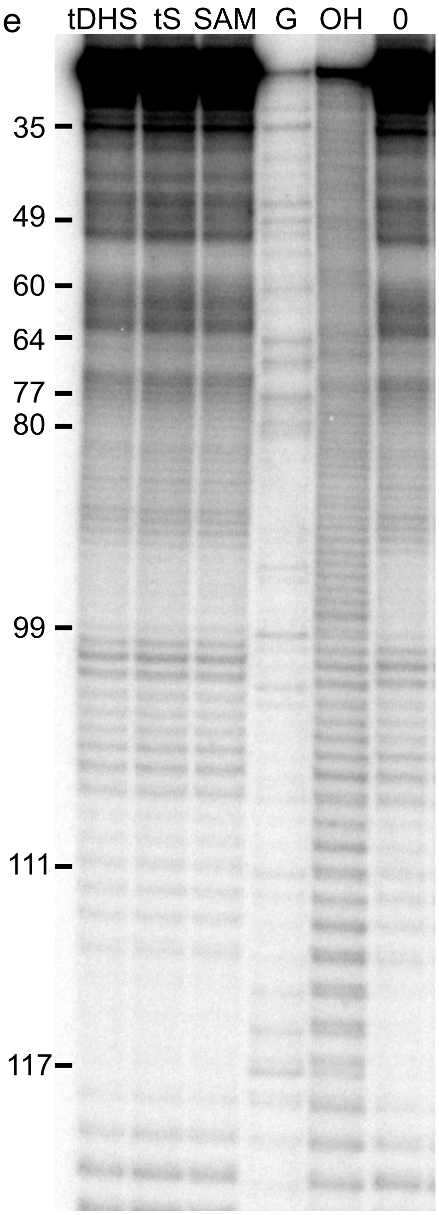
tDHS
tS
SAM
G
OH
0
35
49
60
64
77
80
99
111
117

f

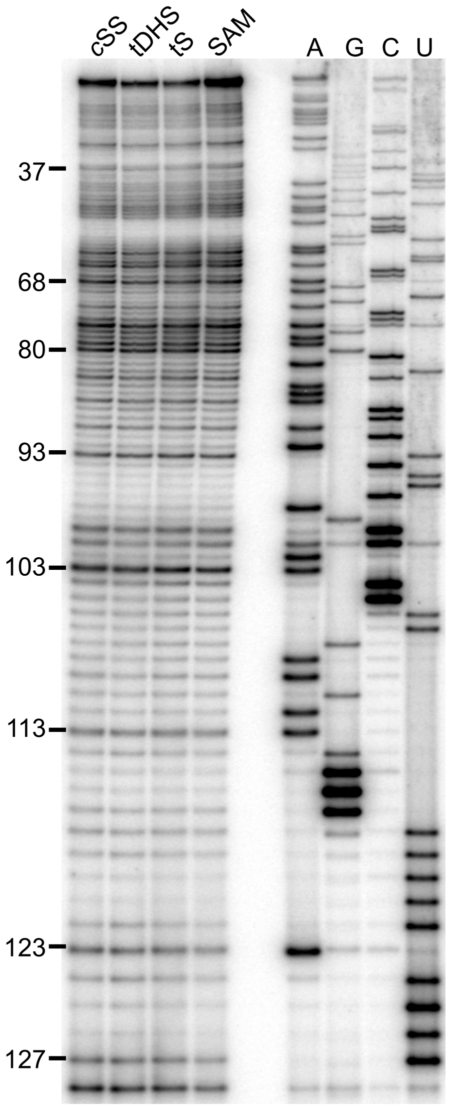
cSS
tDHS
tS
SAM
A
G
C
U
37
68
80
93
103
113
123
127

g

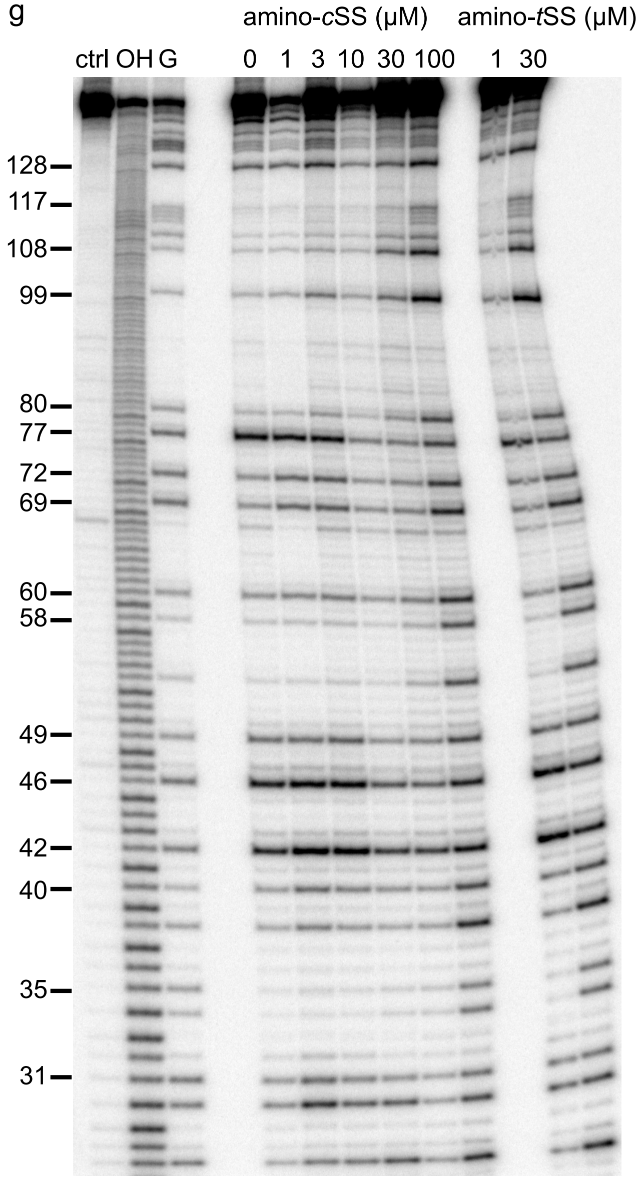
amino-cSS (μM)
amino-tSS (μM)
ctrl
OH
G
0
1
3
10
30
100
1
30
128
117
108
99
80
77
72
69
60
58
49
46
42
40
35
31

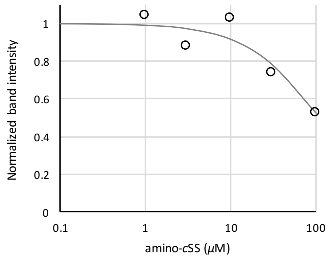
Normalized band intensity
1
0.8
0.6
0.4
0.2
0
0.1
1
10
100
amino-cSS (μM)

**Supplementary Figure 4 | Structural probing of Werewolf-1. a**, SHAPE analysis of Were-1. Left lanes show reverse transcriptase (RT) stops due to ddA ("U") and ddC ("G") incorporation for sequence reference, followed by a no-acylation RT control (ctrl) lane, and SHAPE reactions with amino-*t*SS at concentrations indicated above the gel image. **b,** S1 digestion of Were-1. Left lanes contain a control with no RNA (ctrl), RNase T1 digestion for sequence reference, and a partial hydrolysis lane (OH). The right lanes show S1 digestion in the presence of increasing amino-*t*SS at concentrations indicated above the gel image. Analysis of the band intensity change at positions A44-G46 revealed an apparent $K_D$ value of 0.4 $\mu$M (below). **c,** Terbium (III) footprinting of Were-1 in the presence of increasing amino-*t*SS. Left lanes show RNase T1 digestion for sequence reference, and partial hydrolysis (OH). Middle lanes show $Tb^{3+}$ footprinting in the presence of increasing amino-*t*SS at concentrations indicated above the gel image. Right lanes show control reactions with Were-1 in the presence of 30 $\mu$M controls, as labeled above the gel image. Right, a $K_D$ value of 4.8 μM was calculated based on the change in intensity with increasing amino-*t*SS at nucleotides A113 and U107. **d,** T1 digestion controls. Left lanes contain a control with no RNA, partial hydrolysis (OH), and RNase T1 digestion for sequence reference. Right lanes indicate no ligand (0), followed by Were-1 in the presence of 30 $\mu$M controls, as labeled above the gel image. **e,** In-line probing of Were-1 with 30 $\mu$M controls, as labeled above the gel image, followed by RNase T1 digestion for sequence reference, partial hydrolysis (OH), and a no ligand control (0). **f,** SHAPE of Were-1 in presence of control small molecules. Left lanes show RT stops for Were-1 in the presence of 30 μM amino-*c*SS, tDHS, tS, and SAM. Right lanes show RT stops due to ddT ("A"), ddC ("G"), ddG ("C"), and ddA ("U") incorporation for sequence reference. **g,** T1 digestion of Were-1 in the presence of increasing amino-*c*SS. Left lanes show undigested RNA, partial hydrolysis, and a G-specific sequencing lane. Middle lanes show Were-1 digestion in the presence of increasing amino-*c*SS and the right lane show amino-*t*SS experiments at low and high concentrations, for direct comparison of the ligand-induced structural probing. At high concentration (100 μM) amino-*c*SS mimics the bound profile of amino-*t*SS. The apparent $K_D$ calculation for *c*SS is shown below the gel image and in Fig. 1b, revealing a $K_D$ of 108 $\mu$M.

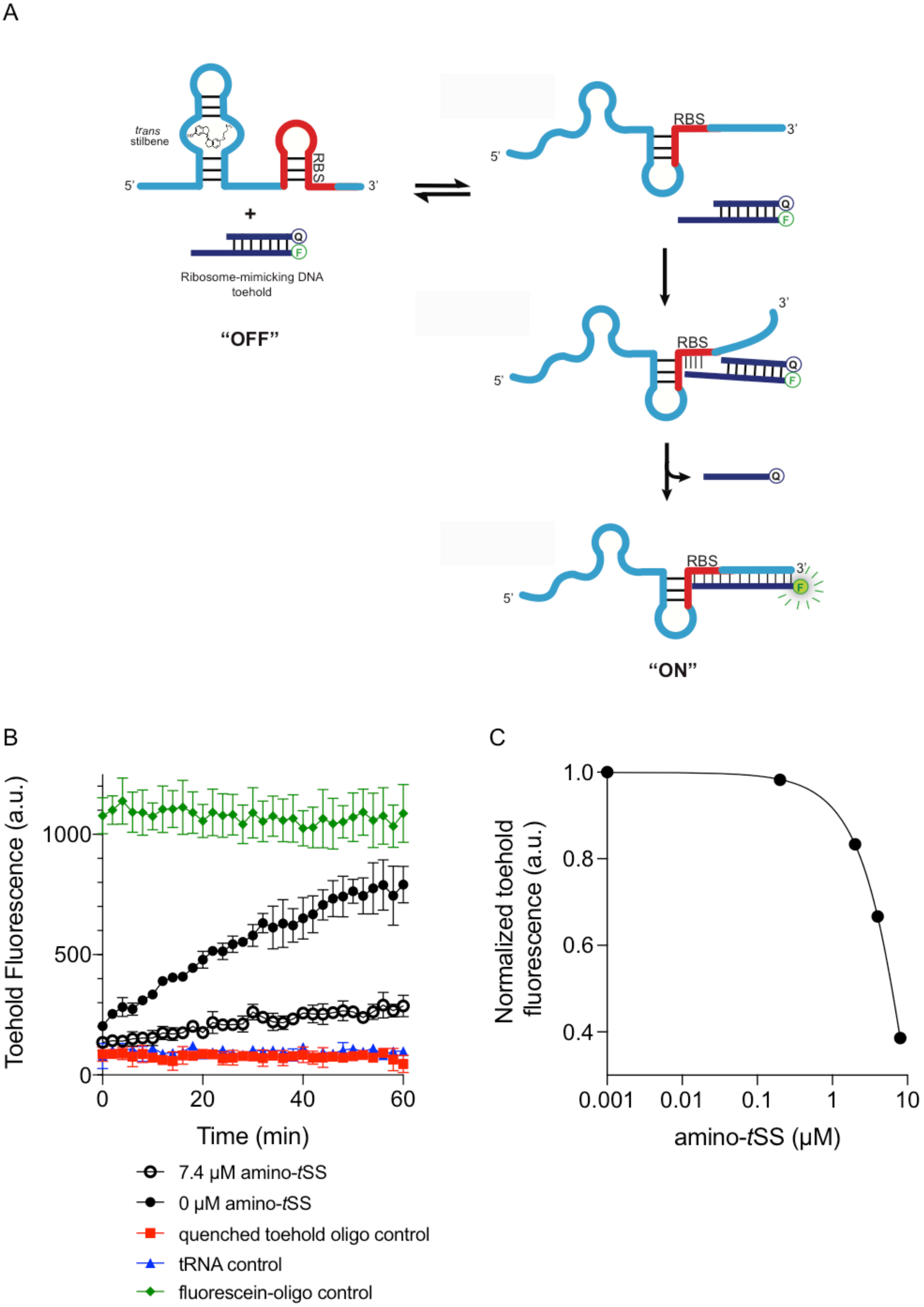
A
trans
stilbene
5'
RBS
3'
+
Q
F
Ribosome-mimicking DNA
toehold
"OFF"
5'
RBS
3'
Q
F
3'
RBS
5'
Q
F
Q
RBS
3'
5'
F
"ON"
B
Toehold Fluorescence (a.u.)
1000
500
0
0
20
40
60
Time (min)
7.4 μM amino-tSS
0 μM amino-tSS
quenched toehold oligo control
tRNA control
fluorescein-oligo control
C
Normalized toehold
fluorescence (a.u.)
1.0
0.8
0.6
0.4
0.001
0.01
0.1
1
10
amino-tSS (μM)

**Supplementary Figure 5 | Binding PAGE-purified Were-1 RNA to a ribosome-mimic. a**, Schematic of strand displacement, where a DNA duplex containing a toehold complementary to the Shine-Dalgarno sequence of Were-1 is displaced when no amino-*t*SS is present, producing a fluorescence. **b**, The presence of amino-*t*SS prevents the quencher strand release, suppressing toehold fluorescence (± SD) over time. Control reactions were performed using the unquenched fluorophore oligo (green) and the quenched toehold oligo (red) for fully quenched fluorescence to provide a window for maximum fluorescence, and a tRNA (blue) lacking the Shine-Dalgarno sequence as a negative control. **c**, Dose-response curve of PAGE-purified Were-1 RNA in the presence of increasing concentrations of amino-*t*SS and analyzed by the toehold-fluorescence assay. Half maximum fluorescence is observed at 6.3 µM amino-*t*SS.

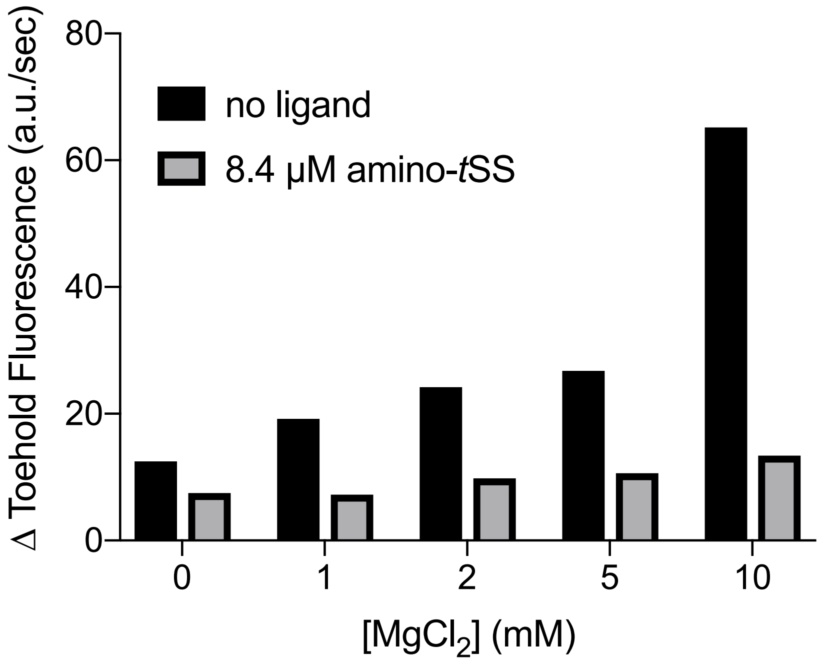


**Supplementary Figure 6** | Co-transcriptional binding of a ribosome-mimic *in vitro* under various $Mg^{2+}$ conditions. Whereas the rate of transcription increases with magnesium (black), the bound state (gray) does not change significantly, suggesting that co-transcriptional binding is minimally affected by magnesium concentration.

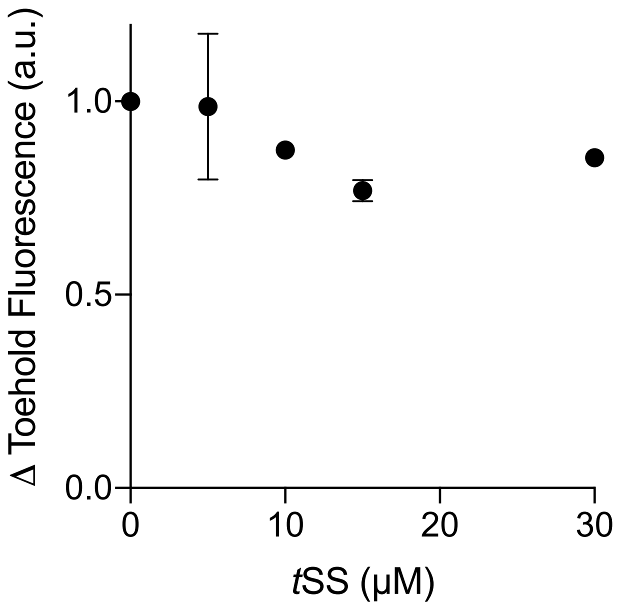


**Supplementary Figure 7 | Co-transcriptional Were-1 binding of a ribosome-mimic *in vitro* in the presence of *t*SS (in 10 % DMSO).** A minor dose-response is observed in the presence of increasing *t*SS concentrations, but full inhibition, as seen for amino-*t*SS, is not observed, possibly due to low solubility of *t*SS.

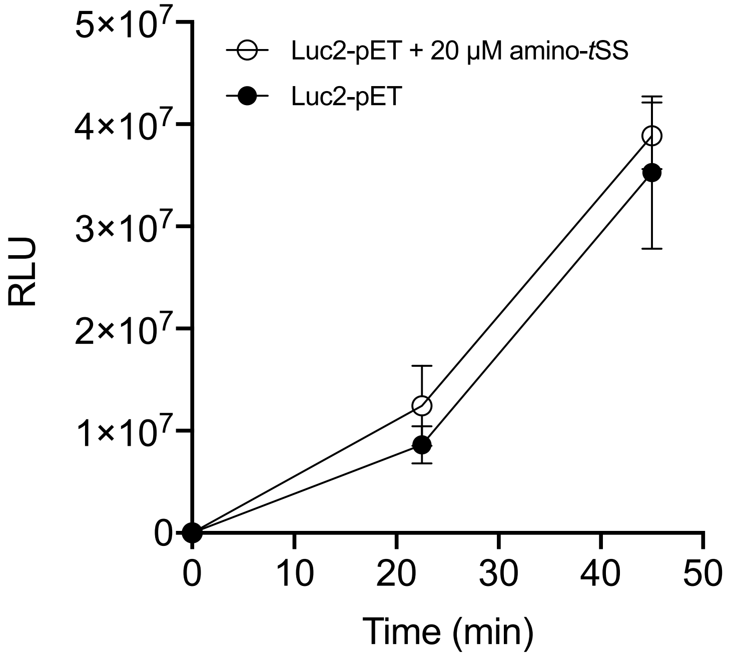


**Supplementary Figure 8 | Translation of a control plasmid, Luc2-pET, lacking the Were-1 riboswitch is not inhibited *in vitro* by amino-*t*SS.** There is no significant difference in Luc2-pET luminescence (± SD) in the presence and absence of amino-*t*SS, suggesting that amino-*t*SS does not inhibit the *in vitro* translation system.

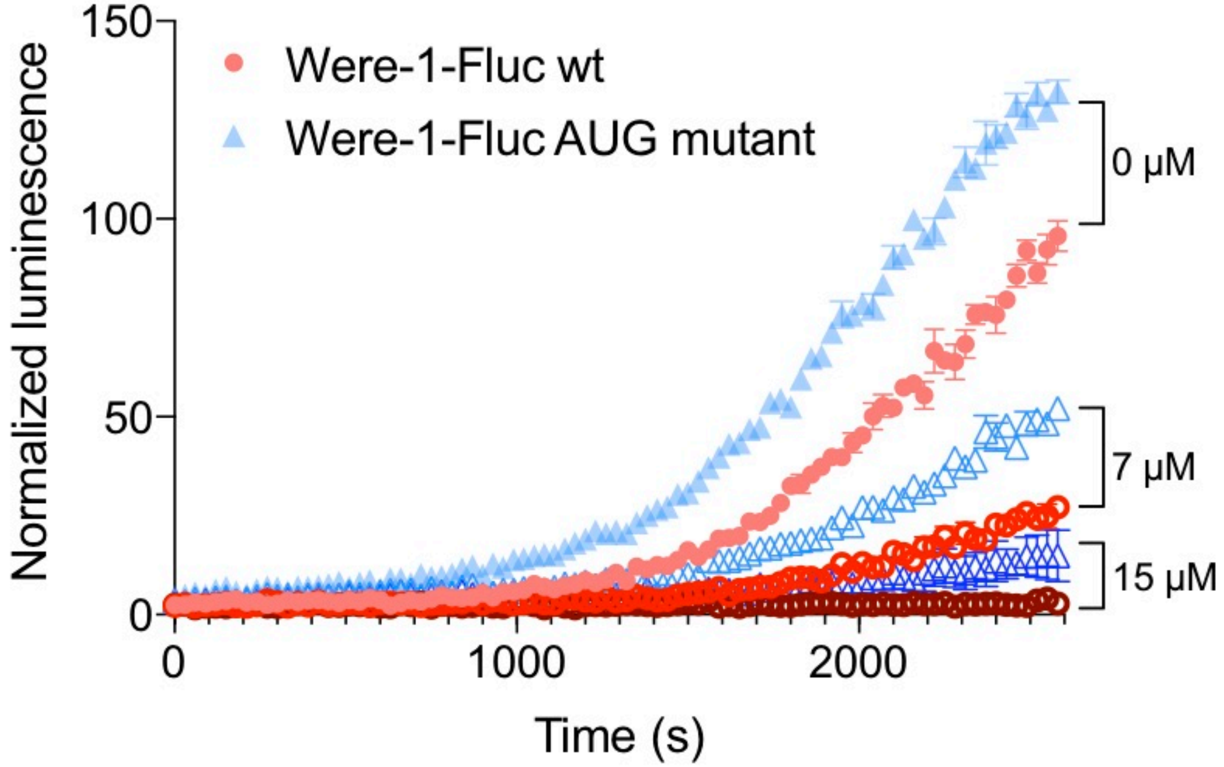


**Supplementary Figure 9 | The effect of the canonical start codon on Were-1-Fluc expression *in vivo*.** Comparison of luciferase expression (± SEM) of Were-1-Fluc wild-type (wt; red) and Were-1-Fluc containing an AUG start codon (Were-1-Fluc AUG mutant; blue), in place of the pool-derived UUG minor start codon, with increasing amino-*t*SS concentrations. Bioluminescence was overall higher in the Were-1-Fluc AUG mutants for all conditions but retained the wt dose-dependent response.

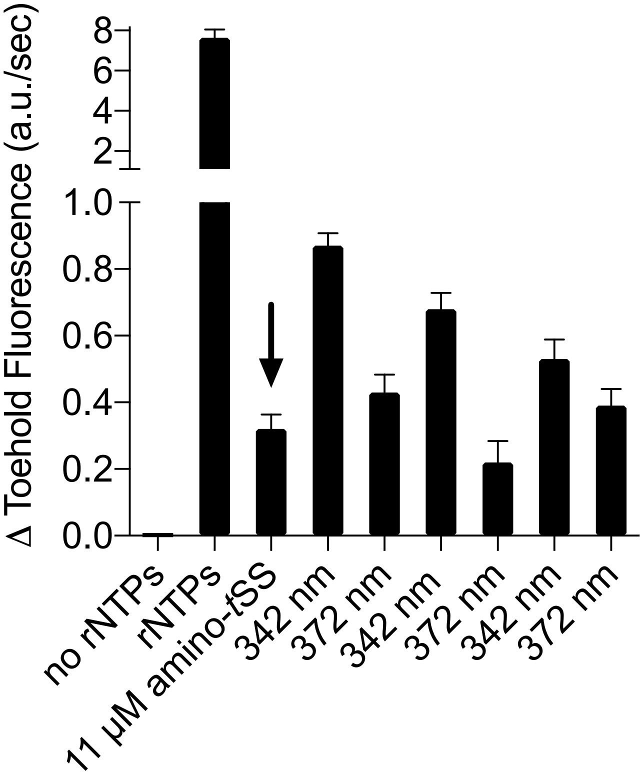


**Supplementary Figure 10 | Co-transcriptional binding of a ribosome-mimic *in vitro* under amino-*t*SS photoisomerization conditions.** Initial transcription of Were-1 without ligand was induced by adding ribonucleoside triphosphates (rNTPs) to a transcription mix, producing robust toehold fluorescence increase (see Fig. 2b for examples of co-transcriptional toehold binding). Upon adding 11 $\mu$M amino-*t*SS (arrow), fluorescence growth decreased, suggesting that Were-1 is bound to the ligand and prevents toehold binding. The sample was then exposed to 342 nm ($\Phi_q$ = $6.8*10^{-5}$ W/cm$^2$) for 60 s, causing photoisomerization of the ligand, and resulting in increased toehold fluorescence slope. Were-1 transcription reaction was subsequently exposed to 372 nm light ($\Phi_q = 1*10^{-4}$ W/cm$^2$) for 60 s, to switch the ligand from *cis* to *trans* conformation, which resulted in decrease of toehold fluorescence slope. This was repeated two more times, as indicated, until the transcription reaction plateaued, and all toehold was expended. The data suggest that the system can be regulated multiple times in one reaction. Error bars represent fluorescence slope error.

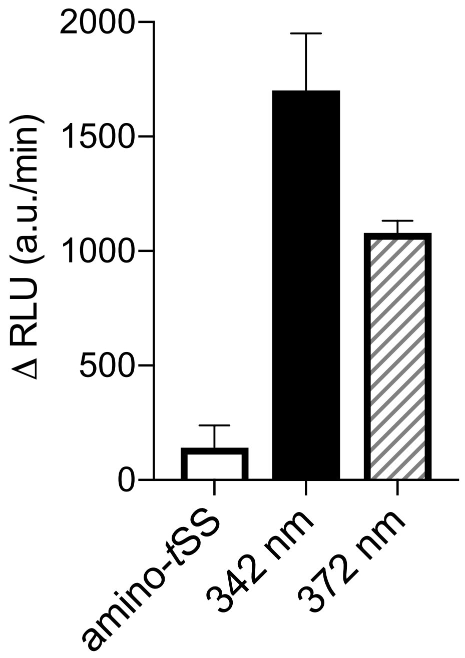


**Supplementary Figure 11 | Were-1 regulates translation *in vitro*.** *In vitro* transcription-translation reaction of Were-1-Fluc mRNA in presence of amino-*t*SS (left column) was irradiated at 342 nm for 60 s, resulting in an increase in luciferase production (± SEM). The reaction was then irradiated at 372 nm for 60 s, resulting in a decrease in luciferase expression rate (± SEM) over time.

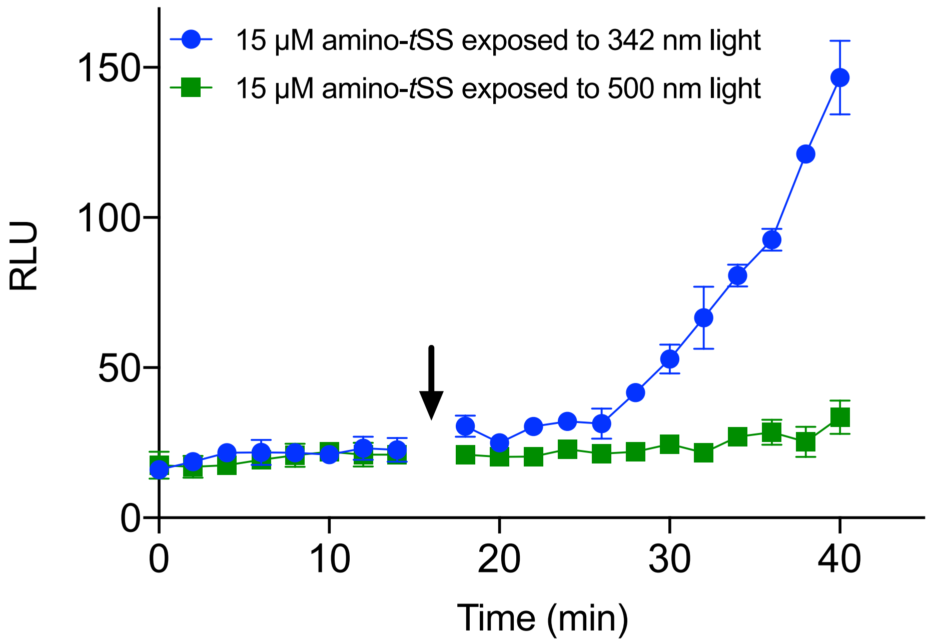


**Supplementary Figure 12 | Were-1 regulates protein expression *in vivo.*** Were-1 *E. coli* bioluminescence (± SEM) was measured in the presence of amino-*t*SS. Initial Fluc expression showed identical increases in bioluminescence for all samples. In the presence of 15 $\mu$M amino-*t*SS, samples were either exposed to (arrow) 342 nm light (blue) to isomerize amino-*t*SS, or 500 nm (green), a wavelength that does not affect amino-*t*SS isomerization. 342-nm–exposed samples showed significantly higher bioluminescence compared to those exposed to 500 nm light.

**Supplementary Table 1 | DNA sequences used for *in vitro* and *in vivo* analysis of the Were-1 riboswitch.**

| | |
|---|---|
| Toehold-Fluorophore Reporter | Rep F: 5'– 5FluorT/TA CCTGCAAGCTTCCCTTTTCAAAATAAAAACCCCT –3'<br>Rep Q: 5'– ATTTTGAAAAGGGAAGCTTGCAGGTA/3IABkFQ –3' |
| Forward Overhang Primer (AL2909) | 5'– CCCgaattcTAATACGACTCACTATAGGGATCCGAAAATTGAAATAGAC –3' |
| Reverse Overhang Primer (AL2912) | 5'– atggcgccgggcctttctttatgtttttggcgtcttCCCTGCAAGCTTCCCTTTTCAAAATAAAAAC –3' |
| Forward Werewolf Primer (AL2881) | 5'– GGGATCCGAAAATTGAAATAGACTCC –3' |
| Reverse Nested Fluc Primer (AL1649) | 5'– gattctgtgatttgtattcagcccata –3' |
| Pool Forward Primer (AL2045) | 5'– TAATACGACTCACTATAGGGATCCGAAAATTGAAATAGAC –3' |
| Pool Reverse Primer (AL2049) | 5'– TACCTGCAAGCTTCCCTTTTC –3' |
| AUG mutant Forward Primer (AL3267) | 5'– 5Phos/ATGAAAAGGGAAGCTTGCAGGgaagac –3' |
| AUG mutant Reverse Primer (AL3268) | 5'– /5Phos/AA TAA AAA CCC CTT CTT CAA GGT TGG CTG AAG –3' |
| C89G mutant Forward Primer (AL3330) | 5'– /5Phos/GACATCTTCAGCCAACCTTG – 3’ |
| C89G mutant Reverse Primer (AL3331) | 5'– /5Phos/GTTTGATGCTTCTGGTCATCTG –3' |
| G69C mutant Forward Primer (AL3328) | 5'– /5Phos/CATGACCAGAAGCATCAAACCAC –3' |
| G69C mutant Reverse | 5'– /5Phos/TGGTAATTCACGGTGCTTACTGC –3' |

| | |
|---|---|
| Primer (AL3329) | |
| C91G mutant Forward Primer (AL3326) | 5'– /5Phos/GATCTTCAGCCAACCTTGAAGAAGG –3' |
| C91G mutant Reverse Primer (AL3327) | 5'– /5Phos/TGGTTTGATGCTTCTGGTCATCTG –3' |
| C84G/G69C compensatory mutant Forward Primer (AL3384) | 5'– /5Phos/GAAACCACATCTTCAGCCAACCTT–3' |
| C84G/G69C compensatory mutant Reverse Primer (AL3385) | 5'– /5Phos/ATGCTTCTGGTCATGTGGTAATTCAC –3' |
| Were-1 DNA | 5'– GGGATCCGAAAATTGAAATAGACTCCGATGGACGGTTGCGTGTATGCAGTAAGCA CCGTGAATTACCAGATGACCAGAAGCATCAAACCACATCTTCAGCCAACCTTGAA GAAGGGGTTTTATTTTGAAAAGGGAAGCTTGCAGGTA –3' |
| Amino-*t*SS pool | 5'– TAATACGACTCACTATAGGGATCCGAAAATTGAAATAGAC1111111111111111111 1111111111111111111111452554554554342255533332232223111111111GAAGAAGGG GTTTTATTTTGAAAAGGGAAGCTTGCAGGTA–3' |
| Were-1-Fluc construct | 5'– TAATACGACTCACTATAGGGATCCGAAAATTGAAATAGACTCCGATGGACGGTTG CGTGTATGCAGTAAGCACCGTGAATTACCAGATGACCAGAAGCATCAAACCACAT |

| | |
|---|---|
| | CTTCAGCCAACCTTGAAGAAGGGGTTTTTATTTTGAAAAGGGAAGCTTGCAGGgaag<br>acgccaaaaacataaagaaaggcccggcgccattctatccgctggaagatggaaccgctggagagcaactgcataaggctatgaaga<br>gatacgccctggttcctggaacaattgcttttacagatgcacatatcgaggtggacatcacttacgctgagtacttcgaaatgtccgttcggt<br>tggcagaagctatgaaacgatatgggctgaatacaaatcacagaatcgtcgtatgcagtgaaaactctcttcaattctttatgccggtgttg<br>ggcgcgttatttatcggagttgcagttgcgcccgcgaacgacatttataatgaacgtgaattgctcaacagtatgggcatttcgcagcctac<br>cgtggtgttcgtttccaaaaaggggttgcaaaaaattttgaacgtgcaaaaaaagctcccaatcatccaaaaaattattatcatggattctaa<br>aacggattaccagggatttcagtcgatgtacacgttcgtcacatctcatctacctcccggttttaatgaatacgattttgtgccagagtccttc<br>gatagggacaagacaattgcactgatcatgaactcctctggatctactggtctgcctaaaggtgtcgctctgcctcatagaactgcctgcg<br>tgagattctcgcatgccagagatcctatttttggcaatcaaatcattccggatactgcgattttaagtgttgttccattccatcacggttttggaa<br>tgtttactacactcggatatttgatatgtggatttcgagtcgtcttaatgtatagatttgaagaagagctgtttctgaggagccttcaggattac<br>aagattcaaagtgcgctgctggtgccaaccctattctccttcttcgccaaaagcactctgattgacaaatacgatttatctaatttacacgaaa<br>ttgcttctggtggcgctcccctctctaaggaagtcggggaagcggttgccaagaggttccatctgccaggtatcaggcaaggatatggg<br>ctcactgagactacatcagctattctgattacacccgagggggatgataaaccgggcgcggtcggtaaagttgttccatttttgaagcga<br>aggttgtggatctggataccgggaaaacgctgggcgttaatcaaagaggcgaactgtgtgtgagaggtcctatgattatgtccggttatgt<br>aaacaatccggaagcgaccaacgccttgattgacaaggatggatggctacattctggagacatagcttactgggacgaagacgaacac<br>ttcttcatcgttgaccgcctgaagtctctgattaagtacaaaggctatcaggtggctcccgctgaattggaatccatcttgctccaacacccc<br>aacatcttcgacgcaggtgtcgcaggtcttcccgacgatgacgccggtgaacttcccgccgccgttgttgttttggagcacggaaagac<br>gatgacggaaaaagagatcgtggattacgtcgccagtcaagtaacaaccgcgaaaaagttgcgcggaggagttgtgtttgtggacgaa<br>gtaccgaaaggtcttaccggaaaactcgacgcaagaaaaatcagagagatcctcataaaggccaagaagggcggaaagatcgccgt<br>gtaa–3' |

**Abbreviations/Key**

3IABkFQ – 3' Iowa black fluorophore quencher

5FluorT – 5' Fluorescein

1 A:G:C:T=1:1:1:1

2 A:G:C:T=17:1:1:1

3 A:G:C:T=1:17:1:1

4 A:G:C:T=1:1:17:1

5 A:G:C:T=1:1:1:17
